# Novel hepatocyte-like liver organoids recapitulate crucial mature hepatic functions

**DOI:** 10.1101/2024.10.29.620824

**Authors:** A. I. Ardisasmita, I. P. Joore, N. Levy, A. Myszczyszyn, A. Marsee, T. Sinnige, J. Ruiter, S. Ferdinandusse, O. Dudaryeva, E. Gruber, V. Daive, R. Levato, B Spee, M.M.A. Verstegen, L.J.W. van der Laan, E. E. S. Nieuwenhuis, I. F. Schene, S. A. Fuchs

**Affiliations:** Department of Metabolic Diseases, Wilhelmina Children’s Hospital, University Medical Center Utrecht, Lundlaan 6, 3584 EA, Utrecht, The Netherlands; Regenerative Medicine Center Utrecht, Uppsalalaan 8, 3584 CT, Utrecht, The Netherlands; Department of Clinical Sciences, Faculty of Veterinary Medicine, Utrecht University, Utrecht, The Netherlands; Institute for Risk Assessment Sciences, Utrecht University, 3584 CM Utrecht, The Netherlands; Laboratory Genetic Metabolic Diseases, Amsterdam UMC, University of Amsterdam, Department of Clinical Chemistry, Amsterdam Gastroenterology Endocrinology Metabolism, Meibergdreef 9, 1105 AZ Amsterdam, The Netherlands; Department of Orthopaedics, University Medical Center Utrecht, Utrecht, The Netherlands; Erasmus MC Transplant Institute, University Medical Center Rotterdam, Department of Surgery, Rotterdam, the Netherlands; Erasmus MC Rare Diseases Center, University Medical Center Rotterdam, Rotterdam, the Netherlands

## Abstract

Accurate liver disease modeling and drug toxicity testing still remain challenging as liver cells *in vitro* poorly resemble adult hepatocytes, as we previously demonstrated using whole transcriptome and cell identity analysis. To address this, we used our insights into hepatic modeling to develop hepatocyte-like liver organoids (HeLLOs), a novel human organoid model with mature hepatocyte functions superior to existing models. HeLLOs are easily established from (small) healthy or diseased liver tissues and rapidly expanded for an extended period in optimized culture conditions. Transcriptomic and functional analyses revealed that differentiated HeLLOs closely resemble fresh primary human hepatocytes (PHHs) and perform key hepatic functions such as gluconeogenesis, drug metabolism, and bile acid synthesis. We developed a HeLLO-based toxicity assay with higher sensitivity in predicting liver toxicity of known liver-toxic drugs compared to the gold-standard PHHs. By modeling disease-related mechanisms, such as bile acid transport, HeLLOs uncover transport-inhibition toxicity mechanisms of known liver toxic drugs. Single cell sequencing analysis of HeLLOs identified a heterogeneous cluster of cells with cholangiocyte-like and hepatocyte-like cells, overall resembling liver regenerative cells. As such, HeLLOs hold great promise for advancing liver disease modeling and drug testing. To our knowledge, HeLLOs are the best expandable liver model for predicting adverse drug reactions as well as modeling various liver disease mechanisms.

## Introduction

The liver is a vital organ that performs numerous essential functions of metabolism, immunity, digestion and detoxification, most of which are carried out by hepatocytes that constitute 80% of the liver volume. Dysfunction in liver-specific pathways within hepatocytes is often linked to diseases, with drug-induced liver injury (DILI) being one of the most significant hepatocyte-related conditions (Andrade, 2019). DILI can manifest as hepatitis, cholestasis or a mix of liver injuries. Several drug-driven disease mechanisms underlie DILI, such as the build up of lipids (steatosis) and the inhibition of transport causing a build up of bile acids (cholestasis). DILI frequently leads to failure of drugs during and after clinical trials, causing delays and increasing the costs of drug development (Dirven, 2021). To predict DILI in the drug development pipeline and to model various liver diseases, many *in vitro* models have been designed to replicate hepatocyte functions. However, most of the models do not proliferate and/or fail to maintain hepatocyte functions for a significant amount of time. These limitations result in inadequate recapitulation of hepatocyte functions and a consequential lack of assay predictivity, thereby hindering drug development, studies of disease mechanisms, and advancements in personalized medicine (Weaver, 2020).

While hepatocytes are challenging to culture, cholangiocytes are highly proliferative *in vitro* (Huch, 2015; Sampaziotis, 2017; Roos, 2022). Previously, cholangiocytes in the mouse and human liver have been shown to transdifferentiate into hepatocytes under certain liver damage conditions *in vivo* (Pu, 2022; Gribben, 2024). As such, others have attempted to use cholangiocytes as an alternative source of hepatocytes *in vitro*. Intrahepatic cholangiocyte organoids (ICO) are established from a cholangiocyte population, are highly proliferative and can be established from small pieces of liver tissue (Huch, 2015; Marsee, 2021). This enables the generation of patient-specific cultures, which are conducive of personalized modeling approaches. Upon differentiation, ICOs have been suggested to recapitulate important hepatocyte functions. However, through comprehensive analysis of various hepatocyte-like cells, we have shown that ICOs still severely lack crucial hepatocyte functions and predominantly resemble cholangiocytes (Ardisasmita, 2022). Thus, ICOs lack important properties to be used effectively for modeling hepatocyte disease and predicting drug toxicity.

Here, we present hepatocyte-like liver organoids (HeLLOs), a highly proliferative human liver model resembling cholangiocytes during expansion, which can be differentiated to express important hepatocyte functions. We show that HeLLOs more closely resemble mature hepatocytes at the transcriptomic and functional level than other hepatocyte models. More specifically, functional analyses show that HeLLOs acquire mature drug metabolism, fatty acid metabolism, bile acid transport, and liver protein secretion functions. Furthermore, these capabilities allow HeLLOs to serve as a reliable platform for modeling and predicting the effects of liver-toxic drugs *in vitro* in terms of cytotoxicity and other toxicity mechanisms. Finally, we show that transferring ICO cultures to HeLLO medium greatly enhances functional hepatic differentiation. Taken together, HeLLOs provide a novel breakthrough technology for patient-derived or healthy liver modeling.

## Results

### HeLLOs are easily established, expanded and more hepatocyte-like than existing liver organoid models

Various hepatocyte-like cells (HLCs) have been published, each with strengths and limitations (Xiang, 2019; Zhang, 2018; Du, 2014; Fu, 2019; Chen, 2018). A significant limitation of the HLCs with strong hepatic phenotypes is their inability to proliferate, suggesting that proliferation and differentiation signals counteract each other. Based on the hypothesis that high proliferation signals hamper the acquisition of mature hepatic phenotype, we defined growth factor-reduced expansion and growth factor-depleted differentiation media for organoid culture (Figure 1A). With these media, we established hepatocyte-like liver organoids (HeLLOs) by mincing small liver tissue pieces or biopsies derived from healthy donors or patients and culturing in HeLLO expansion medium (HeLLO-EM; Figure 1A). HeLLO-EM organoids were highly proliferative and exhibited cystic organoid structures similar to other existing liver-derived organoids (Figure 1B, Figure S1A-B; Huch, 2015; Sampaziotis, 2017). Importantly, HeLLO-EM displayed much higher hepatocyte gene expression than these other organoid models (Figure S1C). Differential gene expression analysis also showed upregulation of genes related to hepatocyte functions in HeLLO-EM (Figure S1D). HeLLOs cultured in hepatic differentiation medium (HeLLO-DM) collapsed into dense organoids and were smaller in size compared to intrahepatic cholangiocyte organoids in differentiation medium (ICO-DM) from Huch et al., which is the liver organoid protocol claiming to derive hepatocytes (Figure 1B). Furthermore, HeLLO-DM displayed much higher expression of important hepatic markers compared to donor-matched ICO-DM (Figure S2A) and remained stable for more than two weeks in culture after differentiation (Figure S2B). Finally, using immunofluorescence staining we confirmed that HeLLO-DM showed stronger protein expression of the important hepatocyte markers ALB, CYP3A4, and CYP2E1, compared to ICO-DM derived from the same donor (Figure 1C).

**Figure 1:**
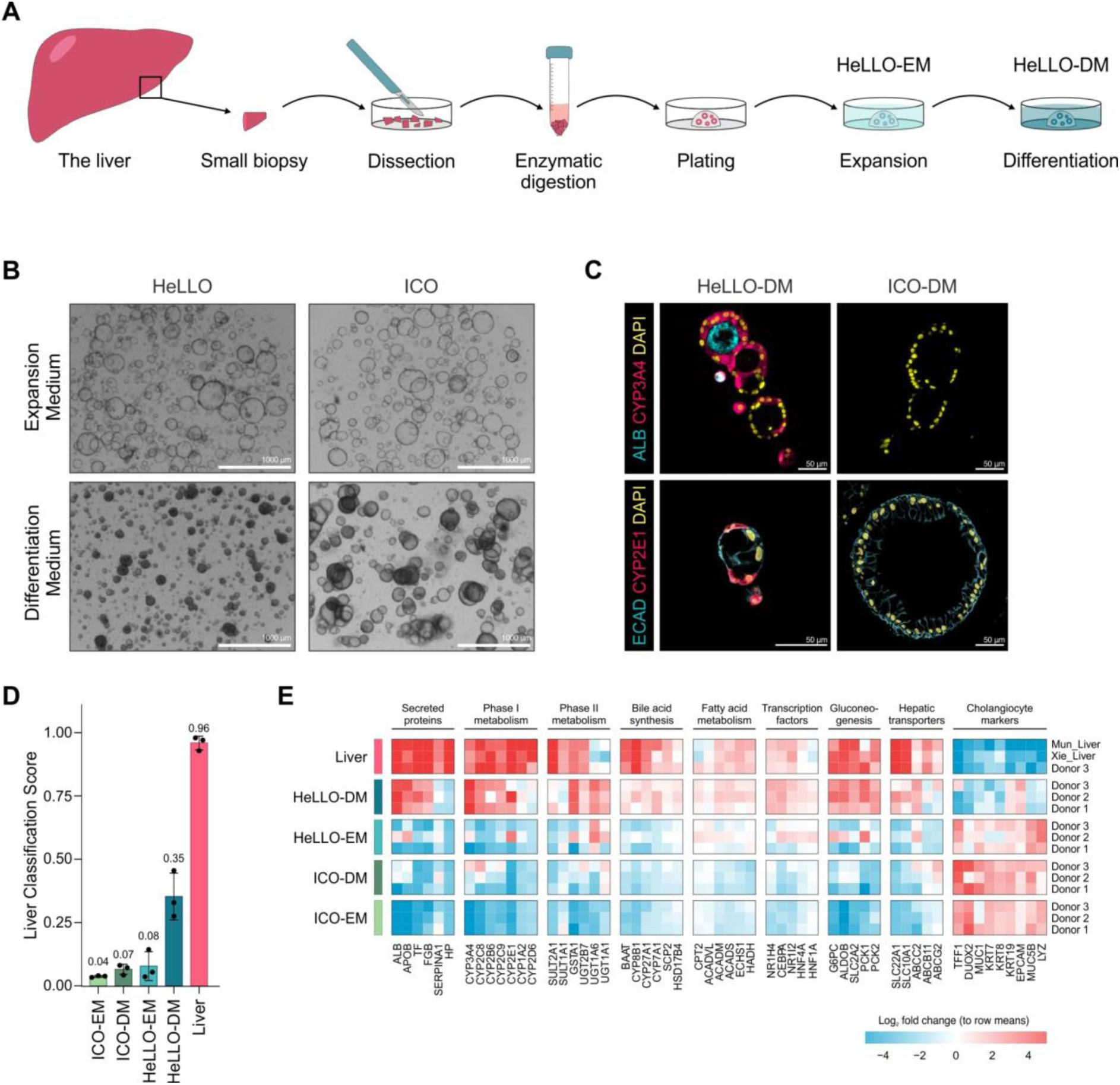
HeLLO establishment, expansion and strong hepatic phenotype. **(A)** Schematic overview of the HeLLO establishment protocol. **(B)** HeLLO brightfield imaging showed similar morphology in the expansion medium (EM) compared to ICOs, while collapsing into distinct dense organoids in the differentiation medium (DM). **(C)** Immunofluorescence staining showed strong expression of liver proteins albumin, CYP3A4, E-cadherin and CYP2E1 in HeLLO-DM but not in ICO-DM (both at differentiation day 8). **(D)** CellNet liver classification of transcriptomic data resulted in a higher liver classification in HeLLO-DM compared to ICO-DM. **(E)** Heatmap analysis of transcriptomic data indicated that HeLLOs gain hepatocyte phenotypes while losing some cholangiocyte markers upon differentiation. Additional liver samples were integrated using HLCompR.

### Removal of growth factors drives hepatocyte differentiation of HeLLOs

To further demonstrate that the removal of growth factors leads to improved hepatic phenotype in organoid culture, we screened the growth factor-rich ICO-DM. Removal of the growth factors in ICO-DM resulted in a sharp increase of hepatocyte marker gene expression levels indicating that removal of growth stimuli is one of the most important factors for hepatic differentiation (Figure S2C). Interestingly, ICOs cultured in ICO-EM and subsequently differentiated in ICO-DM without growth factors (ICO-DM -GF) still expressed lower hepatocyte gene expression levels compared to HeLLOs cultured in HeLLO-DM (Figure S2D), indicating that both HeLLO-EM and HeLLO-DM media are important to obtain better hepatocyte phenotypes compared to ICO-EM and ICO-DM. Possibly, the growth factor removal only partially explains the improved hepatic phenotype as other factors in the HeLLO media might contribute as well.

### HeLLOs display a mature hepatic gene expression profile

We performed a whole transcriptomic analysis as a comprehensive evaluation of the hepatic differentiation status of HeLLO-DM. CellNet classification indicated that the liver classification increased remarkably in HeLLOs (Figure 2D). Additionally, Principal component analysis (PCA) of HeLLO-DM showed a clear transition towards the liver in principal component 1, which captures the largest difference between the samples (Figure S2E). Both CellNet and PCA results still suggested an incomplete acquisition of hepatocyte phenotypes, which we confirmed by assessing specific hepatocyte and cholangiocyte markers (Figure 1E, S2F). We observed that upon differentiation, HeLLOs retained some cholangiocyte markers while gaining liver-level expression of important hepatocyte markers. In contrast, ICOs failed to acquire this hepatocyte phenotype, but retained a general cholangiocyte phenotype (Figure 1D-E). Upon closer inspection of the transcriptomic differences between HeLLO-DM and ICO-DM, we discovered that HeLLO-DM specifically upregulated hepatic genes important for energy and drug metabolism and transport pathways (Figure S2G). Furthermore, HeLLO-DM lost the expression of progenitor markers as opposed to ICO-DM (Figure S2F). Together, these data show that upon differentiation, HeLLOs acquire a hepatic phenotype while retaining some degree of cholangiocyte identity.

**Figure 2:**
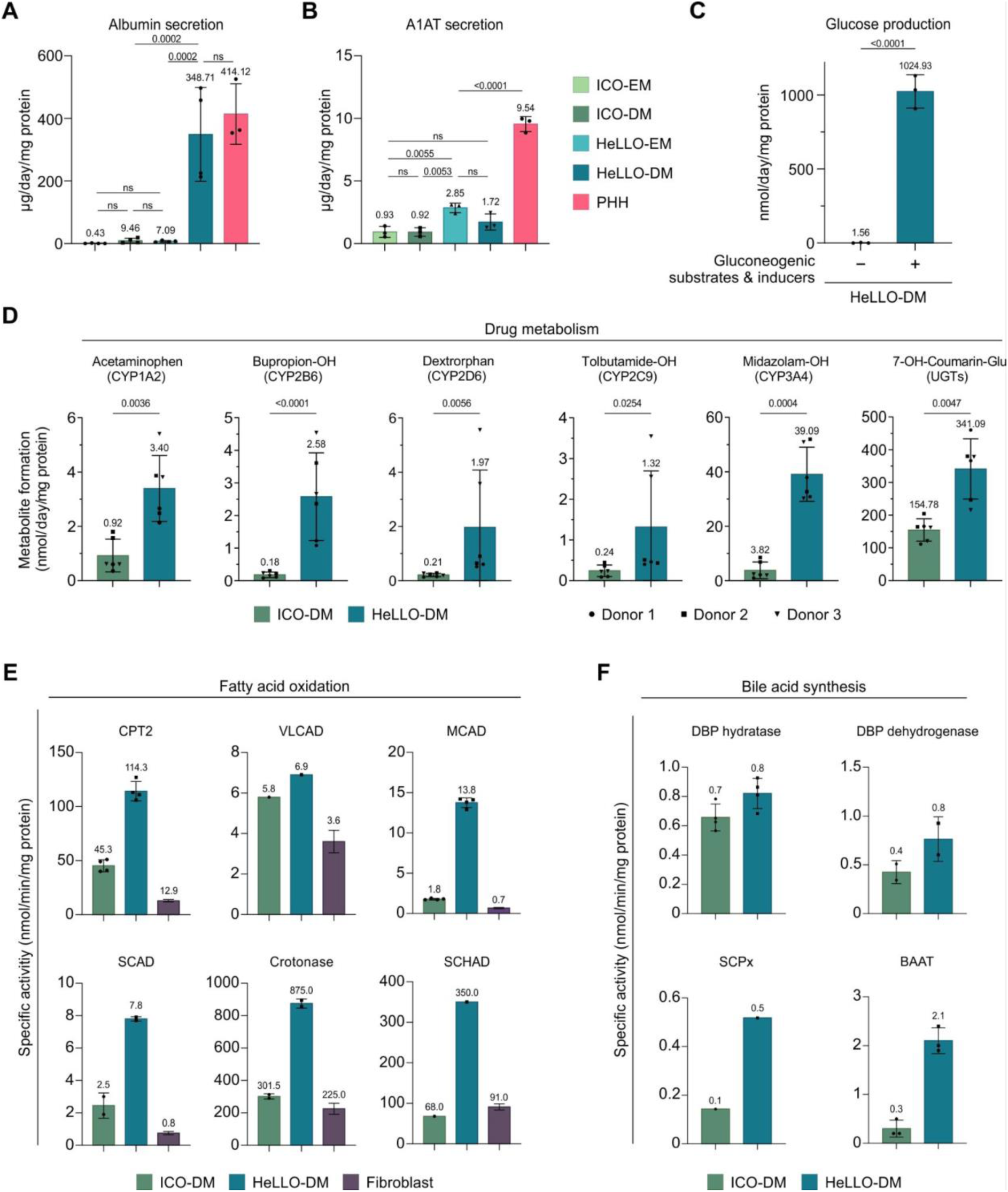
Important liver functions of HeLLOs. **(A)** Albumin secretion by HeLLO-DM (at differentiation day 6), ICO-DM (at differentiation day 8) and sandwich-cultured PHHs (at day 5; see also Figure S4A) (data from two matched donors for HeLLOs and ICOs, three biological replicates from one donor for PHHs). **(B)** A1AT secretion by HeLLO-EM/DM (at differentiation day 0 or 8), ICO-EM/DM (at differentiation day 0 or 8) and sandwich-cultured PHHs (at day 3; see also Figure S4B) (data from two matched donors for HeLLOs and ICOs, three biological replicates from one donor for PHHs). **(C)** Glucose measurement in the media after 24 hours incubation with glucose-free differentiation medium (data from three biological replicates from one donor). Gluconeogenic substrates and inducers were composed of 10 mM Dihydroxyacetone, 20 mM Lactic Acid, 10 μM Forskolin, and 100 μM Dibutyryl-cAMP. **(D)** Drug metabolism assays by measuring metabolite formation in the media after exposing HeLLO-DM and ICO-DM to a mix of CYP-specific compounds (data from three matched donors). **(E)** Fatty acid oxidation enzyme activities measured in the cell lysates of HeLLO-DM, ICO-DM, and fibroblasts (data from 1-4 biological replicates using one matched donor for HeLLOs and ICOs, and panels of fibroblast reference data (CPT2 = 9, VLCAD = 151, MCAD = 25, SCAD = 75, Crotonase = 190, SCHAD = 50)). **(F)** Bile acid synthesis enzyme activities measured in the cell lysates of HeLLO-DM and ICO-DM (data from 1-4 biological replicates using one matched donor for HeLLOs and ICOs).

Using cross-study transcriptome comparison analysis (HLCompR; Ardisasmita, 2022), we benchmarked HeLLOs to other HLCs. In terms of overall resemblance to primary human hepatocytes (PHH), HeLLO-DM was not the most similar HLC-type to PHHs or the liver (Figure S3A & B). However, when considering specific hepatic function gene sets such as gluconeogenesis, drug metabolism, fatty acid metabolism, and bile synthesis, HeLLO-DM outperformed most other HLCs and closely resembled the PHHs (Figure S3C). This shows that HeLLOs strongly express pathways related to specific important hepatic functions, but do not acquire the complete hepatic phenotype.

### HeLLOs exhibit mature hepatocyte functions

We then set out to assess whether HeLLO transcriptomic resemblance to the liver translated to functional activity for these pathways. HeLLO-DM produced albumin at similar levels to sandwich-cultured PHHs while donor-matched ICO-DM produced only low levels of albumin (Figure 2A & S4A). Similarly, HeLLOs demonstrated a significantly higher alpha-1 antitrypsin production compared to ICOs (Figure 2B, Figure S4B). To investigate whether the high gluconeogenic gene expression level in HeLLO-DM also leads to functional gluconeogenesis, we first starved the organoids by exposing them to glucose-free media for 4 hours (Figure S4C) followed by incubation in glucose-free media either with or without gluconeogenic substrates and inducers for 24 hours. HeLLO-DM produced glucose only when gluconeogenic substrates and inducers were present, indicating functional gluconeogenesis (Figure 2C).

Next, we tested the metabolic activities of the most important phase I and II drug metabolizing enzymes (CYP1A2, CYP2B6, CYP2D6, CYP2C9, CYP3A4 and UGTs; Saravanakumar, 2019). Drug metabolism activities of HeLLO-DM significantly outperformed ICO-DM (Figure 2D & S4D). HeLLO-DM also showed higher enzyme activities for different enzymes involved in fatty acid metabolism (Figure 2E). Importantly, activities of several fatty acid oxidation enzymes in HeLLO-DM were higher than in fibroblasts, which are the current standard for clinical testing (Sim, 2002). Enzymes involved in bile acid synthesis were also more active in HeLLO-DM compared to ICO-DM (Figure 2F).

### HeLLOs can be used as liver disease models

An essential function of hepatocytes is the transport of bile into the bile canaliculi. In HeLLOs, the polarization of the organoids is similar to the organization *in vivo*, with the apical lumen of the organoids representing the bile compartment and the basolateral matrix representing the sinusoidal compartment (Figure 3A). We developed a bile acid transport assay by exposing HeLLO-DM to a fluorescent bile acid for 24 hours followed by fluorescence imaging (Figure 3B). We observed clear transport of the fluorescent bile acid into the lumen of the HeLLOs, indicating the capacity to model hepatocyte bile acid transport functions (Figure 3C). ICO-DM is incapable of such bile acid transport, possibly due to low expression levels of NTCP (Figure 3C, 1E). To determine the transporters responsible for bile acid transport, we exposed the HeLLOs with drugs known to inhibit the bile acid transporters. We observed complete ablation of bile acid transport after exposure to zafirlukast (NTCP inhibition; Donkers, 2017) and rifampicin (BSEP inhibition; Weaver, 2020; Figure 3D), suggesting that bile acid transport is NTCP and BSEP dependent. We also developed a lipid accumulation assay by exposing HeLLO-DM to a high dose of free fatty acids (FFA) for 4-10 days (Figure 3E) and observed a clear accumulation of lipid droplets inside HeLLOs (Figure 3F).

**Figure 3:**
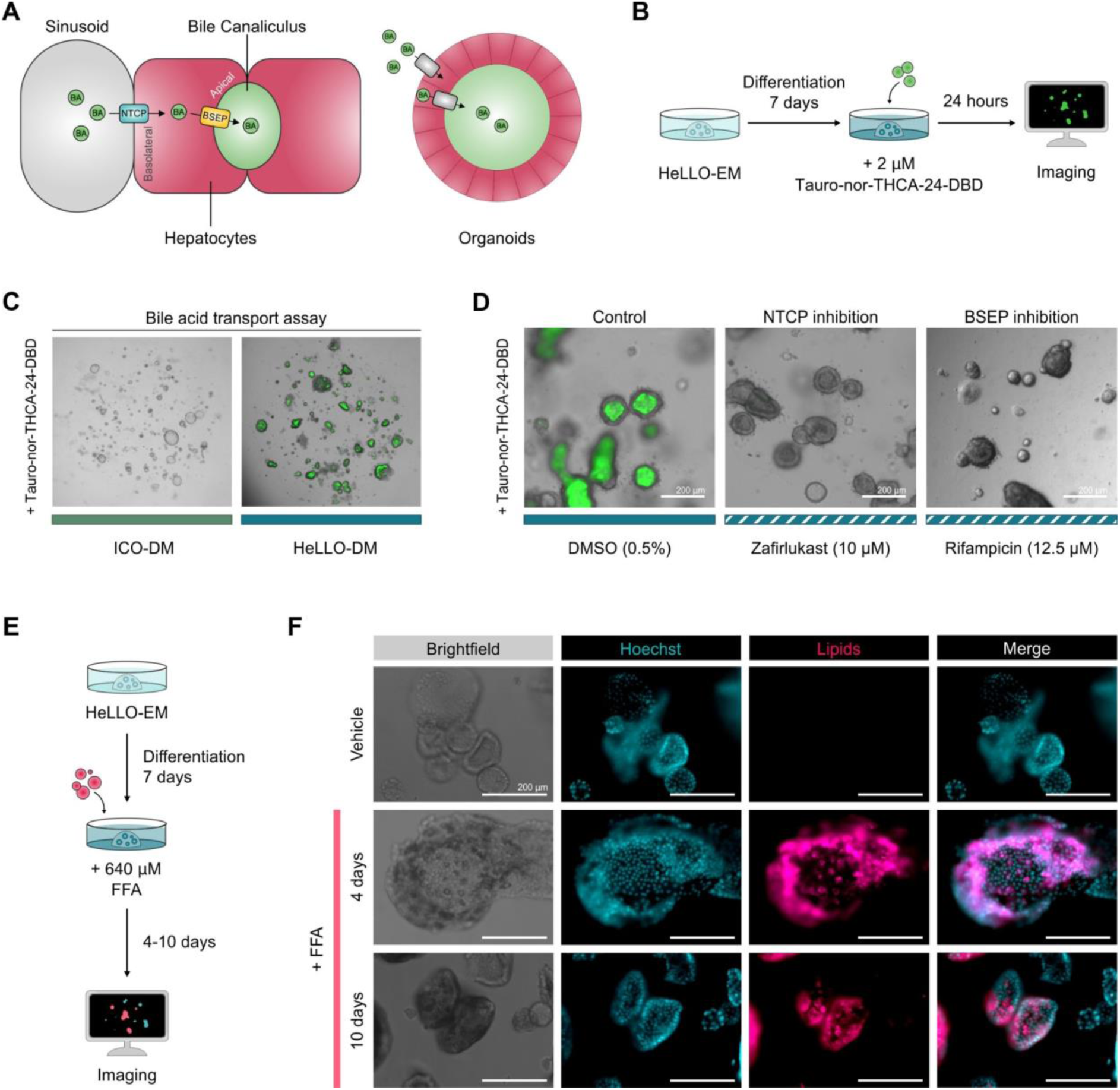
HeLLOs as disease model for liver transport functions and lipid accumulation. **(A)** Illustration of polarization of HeLLOs and the liver architecture. **(B)** Schematic overview of the bile acid transport assay using fluorescent bile acid analog Tauro-nor-THCA-24-DBD. **(C)** Exposure of the fluorescent bile acid for 24 hours showed clear accumulation of fluorescent signal in the lumen of HeLLO-DM but not in ICO-DM. **(D)** NTCP and BSEP inhibition by addition of zafirlukast and rifampicin, respectively, to HeLLO-DM prior to exposure to the fluorescent bile acids showed complete ablation of fluorescent signal. **(E)** Schematic overview of FFA exposure in HeLLOs. **(F)** Representative fluorescence microscopy images of HeLLO-DM long-term exposure to FFA resulted in a buildup of lipids as measured using BODIPY 493/503.

### HeLLOs can predict drug-induced liver injury

Based on the high drug metabolism activity, we set out to test the capacity of HeLLOs to predict drug-induced liver injury (DILI). Cytotoxic effects were measured by exposing HeLLO-DM and ICO-DM for 24 hours with different concentrations of drugs known to induce hepatotoxicity followed by viability assessment using ATP-based assays (Figure 4A). First, we performed a dose response viability experiment using acetaminophen, diclofenac (two DILI-associated drugs), and metformin (a non-hepatotoxic drug) in HeLLO-DM and ICO-DM.

**Figure 4:**
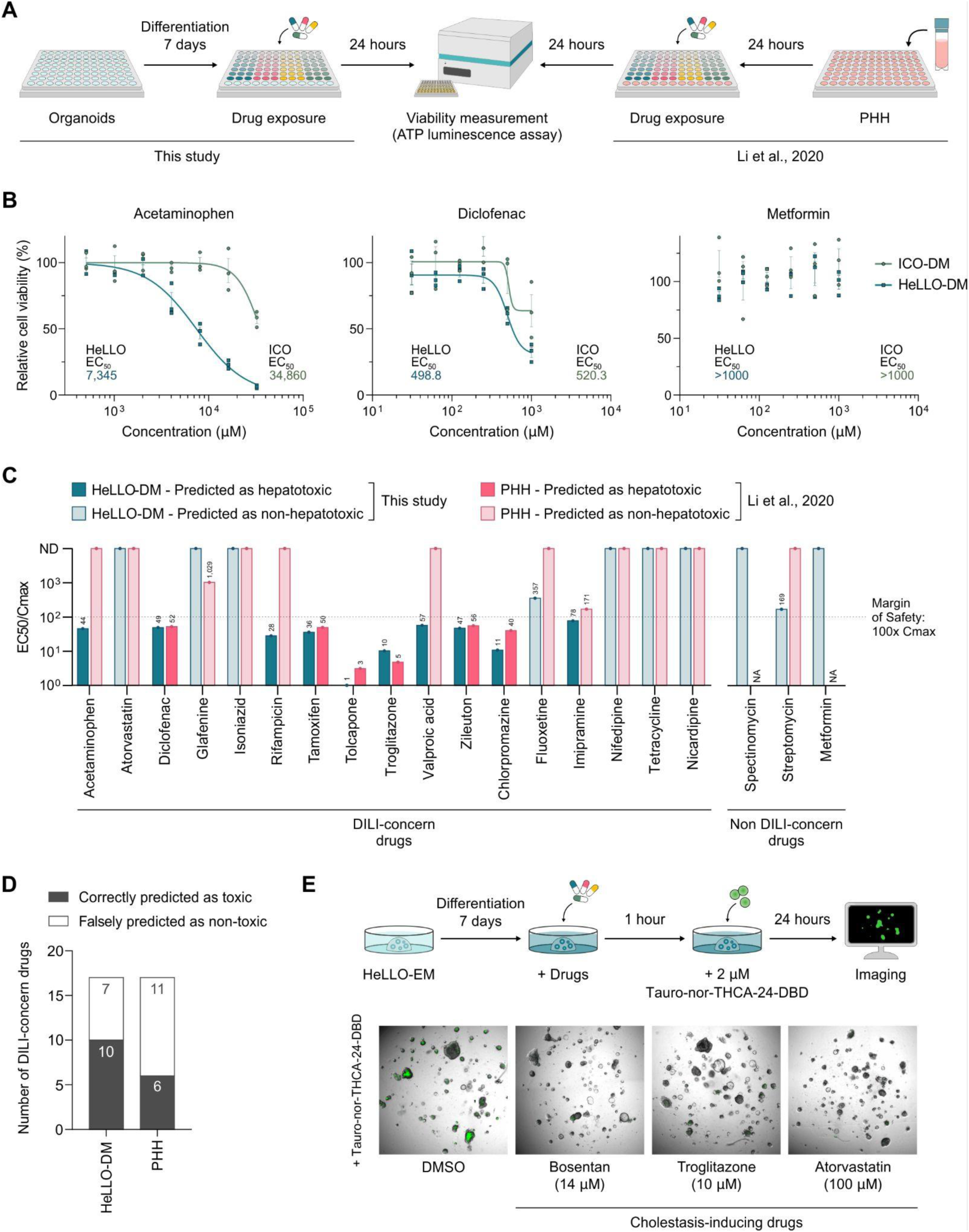
HeLLO toxicity prediction of known liver-toxic drugs. **(A)** Schematic overview of drug exposure on HeLLOs/ICOs and 2D cultured PHHs as performed in Li et al. 2020. **(B)** Comparison of toxicity response to DILI-concern drugs acetaminophen and diclofenac between HeLLOs and ICOs cultured in differentiation media by measuring ATP production as a proxy of cell viability. Metformin was included as a negative control. **(C)** EC50/Cmax results of HeLLOs and 2D cultured PHHs (Li, 2020) after 24-hour exposure to 17 DILI-concern drugs. The dotted line demarcates the 100x Cmax which is used as a margin of safety threshold to assess toxicity (data from one donor, three biological replicates). **(D)** HeLLOs have a higher sensitivity in predicting liver toxicity than 2D cultured PHHs. **(E)** Exposure to known cholestasis-inducing drugs results in a strong decrease in fluorescent bile acid transport in HeLLOs.

HeLLO-DM displayed increased sensitivity to liver toxic drugs compared to ICO-DM, as indicated by the decrease in EC50 (Figure 4B). Next, we compared HeLLO-DM to cultured PHHs, the current gold standard, to predict drug liver toxicity using the data from Li et al. 2020 (Figure 4A, C, Figure S5B). We measured the EC50 of 15 additional DILI-concern drugs (17 in total) and 2 non-DILI-concern drugs (3 in total) in HeLLO-DM (Figure S5A&B). For each drug, we defined the margin of safety (MOS) values as the ratios of EC50 over Cmax (drug concentration in the blood after a therapeutic dose) (Figure 4C). Using an MOS threshold of 100 × Cmax, HeLLO-DM demonstrated greater accuracy in predicting drug liver toxicity than PHHs (Figure 4C). HeLLO-DM correctly predicted toxicity for 10 out of 17 toxic drugs (Figure 4C&D), including the 6 drugs correctly predicted by 2D cultured PHHs and 4 drugs that 2D cultured PHHs incorrectly predicted as safe (Figure 4C).

DILI may not only arise from direct cytotoxicity, but also from cholestasis. To test whether HeLLOs can model drug-induced cholestasis, we exposed HeLLOs to cholestasis-inducing drugs (using a non-cytotoxic concentration) and performed the bile acid transport assay (Figure 4E). We observed a clear absence of bile acid transport after exposure to three known cholestasis-inducing drugs, demonstrating that HeLLOs recapitulate specific *in vivo* toxicity mechanisms of cholestatic drugs (Figure 4E). Together, our data show that HeLLOs are more reliable in predicting DILI than current gold standard and can additionally model underlying drug-related disease mechanisms, such as cytotoxicity and cholestasis.

### Single-cell sequencing of HeLLOs uncovers heterogeneity within HeLLO-DM organoid populations

To assess cellular heterogeneity in HeLLOs, we performed single-cell RNA sequencing on donor-matched HeLLO and ICO samples. After filtering out low quality cells (Figure S6A), we retained 352 ICO-EM cells, 536 ICO-DM cells, 654 HeLLO-EM cells, and 688 HeLLO-DM cells (Figure S6B). Interestingly, each organoid type formed separate clusters, indicating distinct cell identities between the organoid types (Figure 5A). Similar to the bulk RNA sequencing results, the HeLLO-DM cluster displayed the highest hepatocyte marker gene and lowest cholangiocyte marker gene expression levels compared to the other organoid clusters (Figure S6C).

**Figure 5:**
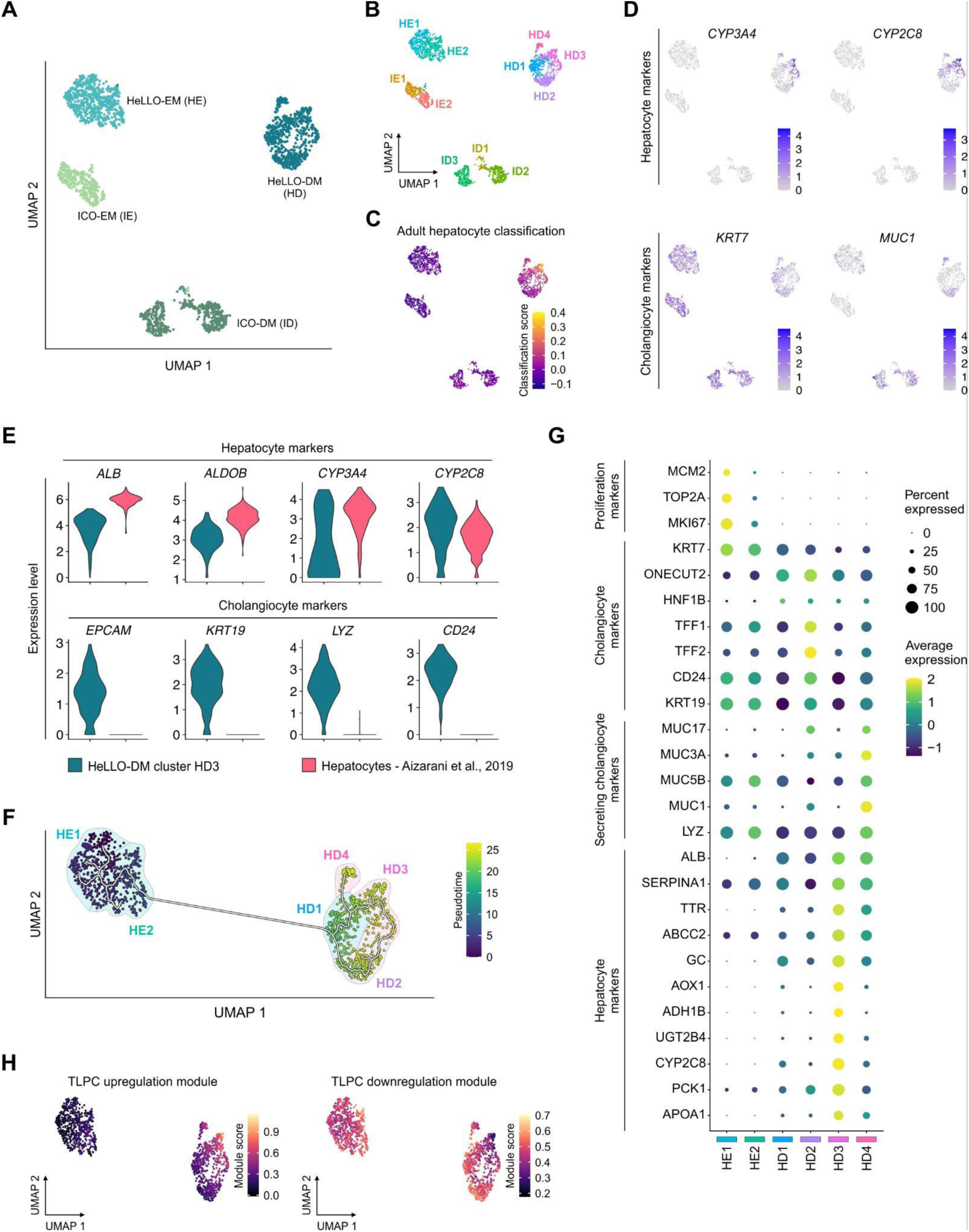
Single-cell sequencing analysis of HeLLOs uncovers heterogeneity within HeLLO-DM organoid populations. **(A)** UMAP visualization of one donor-matched ICO and HeLLO samples in expansion and differentiation media. **(B)** UMAP showing sub clusters of ICO-EM (IE), ICO-DM (ID), HeLLO-EM (HE), and HeLLO-DM (HD) samples. **(C)** UMAP showing SingleR classification scores for adult hepatocytes from the human liver development atlas (Wesley, 2022). **(D)** UMAP showing expression of hepatocyte (top) and cholangiocyte (bottom) marker genes. Color indicates normalized expression data. **(E)** Violin plots of the HeLLO-DM cluster HD3 and hepatocytes from the human liver cell atlas (Aizarani, 2019) showing the normalized expression of hepatocyte (top) and cholangiocyte (bottom) marker genes. **(F)** Trajectory analysis on the HeLLO-EM and HeLLO-DM samples. White line indicates the inferred trajectory, the color bar indicates the pseudotime, and color shading in the background represents an approximation of the clusters. **(G)** Dot plots showing the expression level (color) and percentage of cells (dot size) expressing the proliferation, cholangiocyte, secreting cholangiocyte, and hepatocyte markers on each cluster. This shows clear differences between the clusters with HD3 being the most hepatocyte-like. **(H)** UMAP showing the module score for upregulated and downregulated gene sets in transitional liver progenitor cells (TLPCs) against biliary epithelial cells (BECs) from Pu et al. (2023).

When we further analyzed the clusters, we identified 2 sub clusters of HeLLO-EM (H1 & H2) and ICO-EM (I1 & I2), 3 sub clusters of ICO-DM (ID1-3), and 4 sub clusters of HeLLO-DM (HD1-4) (Figure 5B). Both H1 and I1 clusters highly expressed proliferation markers related to the S and G2M phases (Figure S6D). To identify the cell identity of each cluster, we performed cell type classification using the human liver development atlas as reference (Wesley et al., 2022). Cluster HD3 scored highest in adult hepatocyte resemblance (Figure 5C) and lowest in adult cholangiocyte resemblance compared to the other clusters (Figure S6E), which were classified as adult cholangiocytes (Figure S6E). These were also reflected by the high expression of hepatic marker genes in cluster HD and high expression of cholangiocyte marker genes in the other clusters (Figure 5D). To further compare the HD3 cluster to hepatocytes, we integrated the hepatocyte data from Aizarani et al. (2019) (Figure S6F). Although cluster HD3 expressed similar levels of some hepatocyte marker genes, the cholangiocyte marker gene expression levels were still much higher than hepatocytes (Figure 5E). Additionally, there was no observable liver zonation pattern in the HD3 cluster since both hepatic periportal and pericentral modules were expressed in the same cells although with a tendency towards a periportal phenotype (Figure S6G). Together, this indicates that cells in cluster HD3 are in a biphenotypic state between hepatocytes and cholangiocytes.

To investigate the transcriptomic changes during the HeLLO differentiation process, we performed a trajectory analysis which showed a differentiation trajectory starting from proliferating cluster H1 to non-cycling H2, moving first to cluster HD1 which branches into cluster HD2, HD3, and HD4 (Figure 5F & S6H). When analyzing the differences between clusters (HD1-4), we observed an intermediary cluster lacking hepatocyte and cholangiocyte expression (HD1), a cholangiocyte cluster (HD2), a secreting cholangiocyte cluster (HD4) and a hepatocyte cluster (HD3) (Figure 5G). In this population of cells, HD3 seems to be the end-point according to the trajectory analysis (Figure S6H). Hence, we inferred that the general trajectory is a cholangiocyte to a hepatocyte phenotype, which is reminiscent of a regenerative transdifferentiation ductular response that has been reported in both mice and humans (Pu, 2023; Gribben, 2024). In the mouse study by Pu et al. (2023), liver injury caused by hepatocyte senescence resulted in bile epithelial cells (BECs) transforming into the regenerating biphenotypic transitional liver progenitor cells (TLPCs) which express both cholangiocyte and hepatocyte markers. When scoring our samples using the differentially expressed genes between TLPCs and BECs from Pu et al., the HD3 cluster had the highest score for the gene set upregulated in TLPCs and the lowest score for the gene set downregulated in TLPCs (Figure 5H). This suggests that the HeLLO differentiation process might represent the transdifferentiation found in regenerating BEC ductular response.

### HeLLOs can be established from ICOs

Many biobanks and protocols are built on the ICO protocol. To test whether ICOs can be transformed into HeLLOs, ICOs were grown for four passages, the medium was switched to HeLLO expansion medium in passage five (Switch), grown for several days, and subsequently differentiated using HeLLO differentiation medium (Switch-DM) (Figure S7A). The expression of hepatocyte marker genes of Switch and Switch-DM reached similar levels to HeLLO and HeLLO-DM, respectively (Figure S7B). The overall gene expression of Switch-DM also became more similar to HeLLO-DM than to ICO-DM although still less liver-like than HeLLO-DM (Figure S7C). Switch-DM also transported bile acids effectively (Figure S7D). Culturing in HeLLO-EM medium for multiple passages after switching did not alter the hepatocyte gene expression levels markedly (Figure S7E). Finally, switching directly from ICO expansion medium to HeLLO differentiation medium (Direct-DM), diminished the hepatocyte gene expression levels (Figure S7F-G), once again highlighting the importance of utilizing both optimized expansion and differentiation media to maximize hepatocyte differentiation capacity.

### Potential applications of HeLLOs

HeLLOs possess the potential to be used for both gene and autologous cell therapies to cure liver-related diseases, possibly by providing an engraftable cell source. As a proof of concept, we gene edited HeLLOs using engineered virus-like particles (eVLPs) containing adenosine base editor (ABE) and single guide RNA (sgRNA) targeting the *HBEGF* gene (Figure S8A). Four adenosine bases could be edited in the ABE editing window (A1-A4) (Figure S8B). Addition of the eVLPs to the HeLLOs resulted in high editing efficiency, reaching 80% editing in one of the target bases (Figure S8B).

For cell therapy purposes, HeLLOs have to be grown in compliance with good manufacturing practice. This includes fully-defined components used during culture. However, the typical hydrogels used for organoid culture (BME) originate from animals and the composition is poorly defined. Synthetic hydrogels solve these problems and increase reproducibility and malleability. Therefore, we tested growing HeLLOs in synthetic and well-defined polyethylene glycol (PEG) hydrogels. The hydrogels were additionally supplemented with arginylglycylaspartic acid (RGD) peptides. HeLLOs could proliferate in these PEG hydrogels, although at a slower rate than in BME (Figure S8C). This suggests the necessity of extracellular matrix components to help cell proliferation as previously reported (Ye, 2019).

## Discussion

In this study, we developed a human liver organoid, designated as HeLLOs, that yields novel insights into liver biology and disease mechanisms. HeLLOs can be efficiently established from small liver pieces or biopsies or from ICOs and are readily expanded using optimized media for both expansion and differentiation to enhance hepatocyte characteristics. Differentiated HeLLOs form dense organoids upregulating hepatic expression while losing progenitor and (some) cholangiocyte markers. HeLLO-DM maintained stable hepatocyte gene expression for over two weeks. Functionally, HeLLO-DM produce albumin and alpha-1-antitrypsin at levels comparable to sandwich cultured PHHs and show remarkably high enzymatic activities of several enzymes in bile acid synthesis and fatty acid oxidation. Moreover, to the best of our knowledge, HeLLOs represent the first expandable liver cell model with functional gluconeogenesis and hereby offer the first patient-specific long-term model for studying gluconeogenesis *in vitro*. Furthermore, HeLLO-DM show higher drug enzyme activity of phase I and II drug metabolizing enzymes compared to ICO-DM. In our drug liver toxicity assays, HeLLO-DM more accurately predicted toxicity (HeLLO-DM = 10 out of 17; PHH = 6 out of 17) for known liver toxic drugs than the current gold-standard 2D cultured PHHs and enabled modeling of *in vivo* drug toxicity mechanisms such as cholestasis and steatosis. Single cell sequencing analysis shows cell heterogeneity in HeLLO-DM and presents evidence of the transdifferentiation process from cholangiocytes to hepatocytes. Herewith, we have developed HeLLOs, a human liver organoid platform to model essential liver functions to uncover novel mechanistic insights and develop and test therapies.

HeLLOs resemble cholangiocytes in the expansion phase and gain important hepatocyte functions during differentiation. Cholangiocyte transdifferentiation has been proposed as an *in vivo* regeneration mechanism, in which part of the cholangiocyte population transdifferentiates into hepatocytes in mice and men under certain liver damage conditions (Pu, 2023; Gribben, 2024; Akhurst, 2001; Preisegger, 1999). The ICOs generated using Huch et al. protocols are thought to mimic such damage responses (Huch, 2015). Although the Huch protocol proposes a differentiation method to reach a hepatocyte phenotype, we have shown in this and other studies that ICO-DM hardly recapitulates hepatocyte functions (Ardisasmita, 2022; Lehmann, 2022). In this study, we show that HeLLO-DM fills the gaps of ICO-DM by demonstrating a more mature hepatocyte functionality.

Single-cell sequencing analysis revealed that HeLLO-DM comprises a mixture of cholangiocyte and hybrid cholangiocyte-hepatocyte cells, and result in an incomplete hepatic classification of HeLLO-DM. Possibly, this indicates that the transdifferentiation process in HeLLO-DM is still incomplete. Nevertheless, the heterogeneous cells in HeLLOs may be advantageous for modeling both cholangiocyte and hepatocyte functions. Our comparison with the TLPCs from regenerating livers (Pu, 2023) revealed that HeLLOs resemble the regenerating liver population in terms of downregulated and upregulated genes between TLPCs and BECs. Given the transition from a cholangiocyte phenotype in HeLLO-EM to a hepatocyte phenotype in HeLLO-DM, this possibly indicates that HeLLOs are the first *in vitro* proof of the capacity of human cholangiocyte transdifferentiation towards hepatocytes.

The characteristics of HeLLOs are promising for drug development, particularly for the prediction of drug mechanisms underlying drug-induced liver injury (DILI). Our findings demonstrate that HeLLOs exhibit reliable drug toxicity sensitivity exceeding the current gold-standard PHHs (HeLLO sensitivity: 53%, PHH sensitivity: 27%), and strong functional drug metabolism capacity, showing that HeLLOs are capable of identifying drug toxicity risks. Underlying DILI there are several disease mechanisms leading to toxicity, including cholestasis and steatosis. In the context of cholestasis, we demonstrated proof-of-concept transport inhibition for three drugs known to cause cholestasis. With regards to steatosis, we have already shown that HeLLOs can accumulate lipids upon fatty acid exposure. Furthermore, HeLLOs can generate a diverse panel of donors for population-relevant DILI predictions, which is currently hindered by the availability of donors for PHH-derived and cancer models. Given the FDA’s move from animal testing towards *in vitro* models, HeLLOs offer a compelling alternative to animal testing (Wadman, 2023). Furthermore, it has been proposed that differences between people result in varying toxicological responses that can affect the occurrence of DILI (Chen, 2015). As such, donor diversity represented by HeLLOs could in the future help fill the current gap in liver modeling.

In conclusion, we present HeLLOs as a versatile and promising platform for (personalized) liver disease modeling and drug testing. The high expansion and hepatic differentiation potential underscores their value in addressing the limitations of current hepatocyte-like models and support future advancements.

## Materials and Methods

### Study approval and human subjects

The study was approved by the responsible local ethics committees (Institutional Review Board of the University Medical Center Utrecht (STEM: 10-402/K; TcBio 14-008; Metabolic Biobank: 19–489) and Erasmus MC Medical Ethical Committee (MEC-2014-060)). Tissue biopsies were obtained from donor or explant livers during surgery in the Erasmus MC, Rotterdam. All biopsies were used after written informed consent.

### Organoid establishment and culture

Organoids were established from liver biopsies as previously described (Huch, 2015). Liver biopsies were minced into small pieces and digested by incubation with 2.5 mg/ml Collagenase D (Sigma, 11088866001) in Hanks’ Balanced Salt Solution (details) for 20 min at 37⁰C. The solution was then diluted using wash medium: cold Advanced DMEM/F12 medium (Gibco, 12634028) supplemented with 2 mM GlutaMAX (Gibco, 35050061), 10 mM HEPES (Gibco, 15630080), 100 U/ml Pen-Strep (Gibco, 15140122), and centrifuged at 300 × g, 4⁰C for 5 min. Supernatant was removed and cell pellet was resuspended in a 1:1 mixture of cold Cultrex BME (Bio-Techne, 3536-005-02) and wash medium. The cell suspension was then plated by adding a volume of 30-50 μl droplet into each cell culture well plate. The droplet was incubated at 37⁰C for 15-30 min or until the matrix was solidified.

Hepatocyte-like liver organoids (HeLLOs) were grown by adding HeLLO expansion medium. Hepatic differentiation was done by switching to HeLLO differentiation medium. The medium composition will be made publicly available upon publication in a peer reviewed journal.

To culture intrahepatic cholangiocyte organoids (ICOs), ICO seeding medium was added to the wells for the first 3-7 days after seeding of the biopsies or until organoids were formed (Huch et al., 2015). ICO seeding medium consists of Advanced DMEM/F12 medium (Gibco, 12634028) supplemented with 2 mM GlutaMAX, 10 mM HEPES, 100 U/ml Pen-Strep, 2% B27 without vitamin A (Gibco, 12587010), 10 mM Nicotinamide (Sigma, N0636), 1.25 mM N-Acetylcysteine (Sigma, A9165), 10% RSPO1 conditioned media (homemade), 10 nM Gastrin (Tocris, 3006/1), 50 ng/ml EGF (Peprotech, AF-100-15), 100 ng/ml FGF10 (Peprotech, 100-26), 25 ng/ml HGF (Peprotech, 100-39), 50 μg/ml Primocin (InvivoGen, ant-pm-2), 5 μM A83-01 (Tocris, 2939/10), and 10 μM Forskolin (Tocris, 1099/10), 30% Wnt conditioned media (homemade), 25 ng/ml Noggin (Peprotech, 120-10C), and hES cell cloning recovery solution (Stemgent, 010014500). After organoids were formed, the medium was changed into ICO expansion medium, which was based on ICO seeding medium but without Wnt conditioned media, Noggin, and hES cell cloning recovery solution. Differentiation towards hepatocyte was initiated by culturing the organoids using ICO expansion medium supplemented with 25 ng/ml BMP7 for 5–7 days followed by ICO differentiation medium, consisting of Advanced DMEM/F12 medium supplemented with 2 mM GlutaMAX, 10mM HEPES, 100 U/ml PenStrep, 2% B27 without vitamin A, 1.25mM N-Acetylcysteine, 10 nM Gastrin, 50 ng/ml EGF, 25 ng/ml HGF, 100 ng/ml FGF19 (Peprotech, 100-32), 50 μg/ml, Primocin, 500 nM A83-01, 25 ng/ml BMP7 (Peprotech, 120-03), 10 μM DAPT (Sigma, D5942), and 30 μM Dexamethasone (Sigma, D4902) for 8 days.

ICOs grown according to Sampaziotis et al. (S-ICOs) (Sampaziotis, 2017 and Tysoe, 2019) were established by adding S-ICO expansion medium to the seeded biopsies. S-ICO expansion medium was based on William’s E medium (Gibco, 22551089) supplemented with 2 mM GlutaMAX, 20 mM HEPES, 100 U/ml Pen-Strep, 10 mM Nicotinamide, 10% RSPO1 conditioned media, 20 ng/ml EGF, 50 μg/ml Primocin, 14 mM Glucose (Sigma, G7021), 1x ITS (Gibco, 41400045), 17 mM Sodium Bicarbonate (Sigma, S6014), 6.3 mM Sodium Pyruvate (Gibco, 11360070), 0.2 mM 2-phospho-L-ascorbic acid trisodium salt (Sigma, 49752), 0.1 μM Dexamethasone, and 100 ng/ml DKK-1 (Abcam, ab155623). All organoid media were refreshed every 3-4 days and organoids were passaged 1:4-1:8 every week by either mechanical disruption or enzymatic digestion using TrypLE Express Enzyme (Gibco, 12604013). If organoids were passaged by enzymatic digestion into single cells, 10 μM Y-27632 (Abcam, ab120129) was added to the culture media for 3-4 days or until organoids were formed.

### Primary human hepatocyte culture

Cryopreserved primary human hepatocytes (PHHs; BeCytes Biotechnologies) were thawed and resuspended in hepatocyte plating medium: William’s E medium supplemented with 2 mM GlutaMAX, 1% Non-Essentials Amino Acid Solution (Gibco, 11140050), 100 U/ml PenStrep, 1x ITS+ Premix (Corning, 354352), 1 mM N-acetylcysteine, 100 nM Dexamethasone, and 5% FBS (Gibco, 26400044). The cells were then plated on collagen-coated plates with the recommended density from the supplier. After 6-8 hours, the medium was switched to HPM without FBS and was refreshed daily.

### Immunofluorescence staining

Organoids were isolated from BME using Dispase (STEMCELL Technologies, 07923) for 15 min at 37⁰C and fixated using 4% formaldehyde (Klinipath, 4078-9001) for 20 min at room temperature, washed with PBS, embedded in 2% agarose, transferred into tissue cassettes, dehydrated, and embedded into paraffin blocks. Sections were prepared using a microtome and hydrated. Antigen retrieval was performed by incubating the sections in either Tris-EDTA (pH 9.0) or citrate buffer (pH 6.0) for 30 min at 98⁰C followed by incubation for 30 min at room temperature. Sections were permeabilized using PBS supplemented with 0.1% Triton X-100 and blocked with 10% Normal Goat Serum (Thermo Fisher, 50197Z) for 30 min at room temperature. Sections were then incubated with primary antibodies at 4⁰C overnight, washed, incubated with secondary antibodies for 1 hour at room temperature, counterstained with DAPI, and mounted using FluorSave (Merck, 345789). Antibodies and the dilutions are listed in Supplementary Table 1. Images were acquired using Leica Dmi8 Thunder Imager microscope (Leica) and processed using Leica Application Suite X, Adobe Photoshop 2023, or ImageJ.

### RNA isolation and RT-qPCR

Total RNA was isolated using PureLink™ RNA Mini Kit (Invitrogen, 12183025) according to the manufacturer’s instructions and subsequently stored at -80⁰C until further processing. RNA concentration was determined using Qubit™ RNA Broad Range Assay Kit (Invitrogen, Q10211) and Qubit™ 4 Fluorometer (Invitrogen). Synthesis of cDNA was performed using T100 Thermal Cycler (Bio-Rad) and iScript™ cDNA Synthesis Kit (Bio-Rad, 1708891) according to the manufacturer’s recommendation. Quantitative PCR was performed using 20 ng cDNA, 0.75 μM forward and reverse primers, and iQ™ SYBR^®^ Green Supermix (Bio-Rad, 170-8886) on a CFX96 Touch Real-Time PCR Detection System (Bio-Rad). The PCR cycles consisted of 39 cycles of 95⁰C for 10 sec and 62⁰C for 30 sec. Gene expression was calculated using the ΔΔCt method normalized to the housekeeping gene heterochromatin protein 1 binding protein 3 (*HP1BP3*). Primer sequences are listed in Supplementary Table 2.

### Bulk RNA sequencing and analysis

Messenger RNA was isolated from total RNA using Poly(A) Beads (NEXTflex). RNA integrity was assessed using Agilent RNA 6000 Nano Kit (Agilent, 5067-1511) and RNA concentration was determined using Qubit RNA High Sensitivity Assay Kit (Invitrogen, Q10211). RNA samples with RIN > 8.0 were used for sequencing. Sequencing library was prepared using the Rapid Directional RNA-Seq Kit (NEXTflex) and sequenced on a NextSeq500 (Illumina) to produce 75 base reads (Utrecht Sequencing Facility).

Raw read processing was performed as previously described (Ardisasmita, 2022) using Galaxy web-based platform (https://usegalaxy.eu/) (Afgan, 2018). Reads were assessed using the FastQC tool (Galaxy Version 0.72) and trimmed using Cutadapt (Galaxy Version 1.66.6). Reads were mapped using RNA STAR tool (Galaxy Version 2.7.2b), Gencode human reference genome sequence release 33 (GRCh38.p13), and Gencode comprehensive gene annotation v33 using default parameters. Count matrices were obtained by selecting the “– quantMode GeneCounts” option in the RNA STAR tool.

Counts were normalized by applying the DESeq2 variance-stabilizing transformation (VST) from the “DESeq2” R package (Love, 2014). Differentially expressed genes were identified using the following parameters: “alpha=0.05” and “lfcThreshold=0”. Principal component analysis (PCA) was done using the “ggplot2” R package. Gene expression heatmaps were generated using the “pheatmap” R package and visualized as mean-centered expression per gene. Distance-based similarity score was calculated as previously reported (Ardisasmita, 2022). Enrichment analysis was performed in enrichR (Chen, 2013). CellNet analysis was done using Platform-Agnostic CellNet web application (Lo, 2023).

### Single-cell RNA sequencing and analysis

Organoids were dissociated into single cells using TrypLE Express Enzyme. Viable single cells were determined using DAPI staining and FACS sorted into 384 well plates using BD FACSAria II (BD Biosciences). Each well contains a 50 nl droplet of barcoded primers and 10 µl of mineral oil (Sigma, M8410). After sorting, plates were centrifuged at 2,000 × g for 1 min at 4°C, snap frozen on dry ice, and stored at -80°C until further processing. Single-cell RNA sequencing was performed by Single Cell Discoveries according to an adapted version of the SORT-seq protocol (Muraro, 2016) with primers described in van den Brink et al. (2017). Cells were heat-lysed at 65°C followed by cDNA synthesis. After second-strand cDNA synthesis, all the barcoded material from one plate was pooled into one library and amplified using *in vitro* transcription (IVT). Following amplification, library preparation was done following the CEL-Seq2 protocol (Hashimshony, 2016) to prepare a cDNA library for sequencing using TruSeq small RNA primers (Illumina). The DNA library was paired-end sequenced on an Illumina Nextseq™ 500, high output, with a 1×75 bp Illumina kit (read 1: 26 cycles, index read: 6 cycles, read 2: 60 cycles). Raw read processing was carried out following the CEL-Seq2 protocol (Hashimshony, 2016; https://github.com/yanailab/CEL-Seq-pipeline) which consists of demultiplexing, mapping using Bowtie2 (Langmead, 2012), and read counting using a modified version of htseq-count script (Anders, 2015).

Single-cell RNA sequencing data analysis was carried out using Seurat (v.4.3.0.1; Hao, 2021). Cells that had less than 1000 UMIs (counts) and 400 genes (features) were filtered out from further analysis. Proliferative cells were identified using CellCycleScoring function in Seurat followed by regressing out the difference between G2M and S phase. The dataset was normalized and scaled with SCTransform function. Clusters were determined using FindClusters function with resolution = 1.0. Cell type classification was performed using “SingleR” R package (Aran, 2019) and the liver development atlas (Wesley, 2022) as reference (de.method = “wilcox”). Data integration with the hepatocyte and ICO samples from the human liver cell atlas (Aizarani, 2019) was done using scVI (Gayoso, 2022). Module scores were calculated using the AddModuleScore function from Seurat. The list of genes used for each module can be found in Supplementary Table 3. Trajectory analysis was performed using Monocle3 (Cao, 2019).

### Protein concentration measurement

Organoids were harvested from the BME using Dispase for 15 min at 37⁰C, washed twice with PBS, and lysed with NP-40 lysis buffer (Thermo Fisher, J60766.AK) supplemented with 1x Halt Protease Inhibitor Cocktail (Thermo Fisher, 10516495), 1x Phosphatase Inhibitor Cocktail 2 (Sigma, P5726), and 1x Phosphatase Inhibitor Cocktail 3 (Sigma, P0044). Protein concentration was measured using Pierce BCA Protein Assay Kit (Thermo Fisher, 23225) according to the manufacturer’s recommendation. Absorbance was measured using CLARIOstar Plus multi-mode plate reader (BMG Labtech)

### Albumin and alpha-1-antitrypsin secretion assays

To measure albumin and A1AT secretion, culture medium was refreshed and subsequently collected after 24 hours. The amount of albumin and A1AT in the medium was then determined using Albumin Human ELISA Kit (Invitrogen, EHALB) and Human alpha-1-Antitrypsin ELISA Kit (AssayPro, EA5101-1) respectively.

### Glucose production assay

Organoids were plated on 12 well plates in 6 x 30 µl droplets of BME per well. After 7 days of differentiation, the medium was switched to glucose-free medium: HeLLO differentiation medium but with William’s E Medium without glucose (MyBioSource, MBS653064). To ensure glucose depletion from the medium and BME, glucose-free medium was refreshed every hour 4-5 times. After the last refreshment, the cells were incubated with glucose-free medium either in the presence or absence of gluconeogenic substrates and inducers for 24 hours. Gluconeogenic substrates and inducers were composed of 10 mM Dihydroxyacetone (Sigma, 820482), 20 mM Lactic Acid (Sigma, L7022), 6.3 mM Sodium Pyruvate, 10 μM Forskolin, and 100 μM Dibutyryl-cAMP (Tocris, 1141/10). Glucose in the medium was measured using Amplex Red Glucose/Glucose Oxidase Assay Kit (Invitrogen, A22189) according to the manufacturer’s recommendation.

### CYP activity assay

CYP activity was measured by adapting the previously published protocol (Bouwmeester, 2023). A cocktail of CYP substrates was added to the medium: 5 μM midazolam (Bufa B.V), 20 μM tolbutamide (Sigma, T0891), 20 μM bupropion (Sigma, B102), 15 μM phenacetin (Sigma, 77440), 12 μM 7-hydroxycoumarin (Sigma, H24003), and 15 μM dextromethorphan (Santa Cruz Biotechnology, sc-278927). The cocktail was allowed to incubate for several time points (4, 8, 16, and 24 hours). After which, 400 μl conditioned medium was placed into a glass vial, mixed with 400 μl MeOH (0.1% (v/v) formic acid), and subsequently stored at -20⁰C until further processing.

The following compounds were used as standards for LC-MS/MS analysis: midazolam, 1-hydroxymidazolam (Sigma, UC430), tolbutamide, 4-hydroxytolbutamide (LGC Standards, TRC-H969850), bupropion, hydroxybupropion (Sigma, H-066), phenacetin, acetaminophen (Sigma, A7085), 7-hydroxycoumarin, 7-hydroxycoumarin glucuronide (LGC Standards, TRC-H924880), dextromethorphan, and dextrorphan (Sigma, PHR1974). All standards were prepared in the same matrix as the cell culture medium. The analysis was performed in a single run using a Shimadzu triple-quadrupole LCMS 8050 system with two Nexera XR LC-20AD pumps, a Nexera XR SIL-20AC autosampler, a CTO-20AC column oven, an FCV-20AH2 valve unit (Shimadzu). Both substrates and metabolites were separated on a Synergi Polar-RP column (150 x 2.0 mm, 4 μm, 80 Å) with a 4 x 2 mm C18 guard column (Phenomenex). The mobile phase consisted of 0.1% (v/v) formic acid in Millipore (A) and 0.1% (v/v) formic acid in MeOH (pH 2.7; B), and was set as 100% A (0–1 min), 100% to 5% A (1–8 min), 5% A (8–9 min), 5% to 100% A (9–9.5 min), and 100% A (9.5–12.5 min). The total run time was 12.5 min, and the flow rate was 0.2 ml/min. Peaks were integrated using LabSolutions software.

### Fatty acid oxidation and bile acid synthesis activity assays

Enzymatic activities of enzymes involved in fatty acid oxidation and bile acid synthesis were all measured using the cell homogenates. Cells were homogenized and sonicated in PBS. CPT2 activity was measured using palmitoyl-carnitine (Merck, P1645) as substrate followed by detection of palmitoyl-CoA using ultra high-performance liquid chromatography (UHPLC) on a reversed-phase column (Wanders, 2010). VLCAD activity was measured using C16:0-CoA (Merck, P9716; 0.25 mM) as substrate and ferrocenium hexafluorophosphate (Merck, 388297; 0.4 mM) as electron acceptor, followed by detection of C16:1-CoA and 3-OH-C16-CoA using UHPLC (Wanders, 2010). MCAD activity was measured using 3-phenylpropionyl-CoA (custom synthesis; 0.3 mM) as substrate and ferrocenium hexafluorophosphate as electron acceptor (Merck, 388297; 1.5 mM), followed by detection of 3-phenylpropenonyl-CoA using UHPLC (Wanders, 2010). SCAD activity was measured using butyryl-CoA (Merck, SB 1508; 25 µM) as substrate and ferrocenium hexafluorophosphate as electron acceptor (Merck, 388297; 0.4 mM), followed by detection of crotonyl-CoA and 3-hydroxybutyryl-CoA using UHPLC (Schmidt, 2011). Crotonase (SCEH or ECHS1) activity was measured using crotonyl-CoA (Merck, C6146; 0.34 mM) as substrate followed by detection of 3-hydroxybutyryl-CoA using UHPLC (Peters et al., 2014). SCHAD activity was measured spectrophotometrically using acetoacetyl-CoA (Merck, A1625; 50 µM) as substrate (Wanders, 2010). DBP and SCPx were measured in a single assay using 24-ene-THC-CoA (custom synthesis; 100 µM) as substrate followed by detection of 24-hydroxy-THC-CoA, 24-keto-THC-CoA, and choloyl-CoA using UHPLC (Ferdinandusse, 2000). BAAT activity was measured using taurine (Merck T0625; 20 mM) and choloyl-CoA (custom synthesis; 200 µM) as substrates followed by measurement of tauro-cholate by HPLC-tandem mass spectrometry (Ferdinandusse, 2005).

### Bile acid transport assay

Organoids were plated on 24 well plates in 3 x 10 μl droplets of BME per well and differentiated for 7 days. The medium was then switched to a differentiation medium containing 2 μM Tauro-nor-THCA-24-DBD (GenoMembrane, GM7001). After 24 hours, images were acquired using Leica Dmi8 Thunder Imager microscope.

### Lipid buildup assay

Organoids were plated on 96 well plates in a 5 μl droplet of BME per well and differentiated for 7 days. Lipid buildup was induced by incubating the differentiated organoids with differentiation medium containing 640 μM free fatty acid (FFA) for 4-10 days. The FFA solution was prepared by dissolving Oleic Acid (MedChemExpress, HY-N1446) and Palmitic Acid (MedChemExpress, HY-N0830) separately in 100% ethanol to get 200 mM Oleic Acid and Palmitic Acid solutions. They were then diluted to 8 mM using 10% (w/v) BSA-PBS. Finally, Oleic Acid and Palmitic Acid solutions were mixed with 1:1 ratio to create an 8 mM FFA stock solution (FFA:BSA ≈ 5:1). Medium containing FFA was refreshed every 2-3 days.

For lipid staining, organoids were released from the BME using Dispase for 15 min at 37⁰C, washed twice with PBS, fixated using 4% formaldehyde for 20 min at room temperature, washed twice with PBS, incubated with 1 μg/ml BODIPY 493/503 (MedChemExpress, HY-W090090) and 5 μg/ml Hoechst (Invitrogen, 11534886) for 15 min at 37⁰C, washed twice with PBS, and then moved to a 96 well plate in 100 μl PBS for imaging.

### Drug toxicity assay

Organoids were plated on 96 well plates in a 5 μl droplet of BME per well and differentiated for 7 days. For each drug, seven different concentrations (Supplementary Table 4) were exposed to the culture using Tecan D300e dispenser. For each concentration, three different wells of organoids were exposed and served as biological replicates. Drugs in the form of powder were weighed one day prior the dosing day and stored at -20⁰C. On the dosing day, drugs were dissolved at 200x the maximum dosing concentration as stock solutions to avoid freeze-thaw cycle and to ensure a maximum of 0.5% DMSO concentration for drugs that were dissolved in DMSO. List of the tested drugs and the details can be seen in Supplementary Table 4. Organoids were incubated with the drugs for 24 hours at 37⁰C.

Cell viability was measured using CellTiter-Glo 3D Cell Viability Assay (Promega, G9683) according to the manufacturer’s recommendation. Relative cell viability was calculated using the following formula:

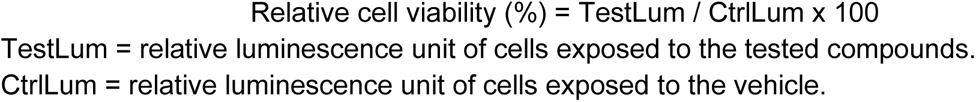

The EC50 of each drug was estimated by fitting the relative cell viability data to a non-linear regression model “variable slope (four parameters)” in GraphPad Prism 9.

### Virus-like particle production and transduction

Engineered virus-like particles (eVLPs) were produced by adopting the previously described protocol (Banskota, 2022). Gesicle Producer 293T cells were plated at a density of 6×10^6^ cells on a T75 flask. After 24 hours, cells were transfected with plasmids expressing VSV-G (4 μg; Addgene #12259), Gag-Pol (3.375 μg; Addgene #35614), Gag-ABE8e (1.125 μg; Addgene

#181751), and gRNA (4.4 μg; Addgene #65777) using Lipofectamine 2000 (Invitrogen, 10696153). The conditioned medium containing eVLPs was then collected 48 hours after transfection, centrifuged at 500 × g for 1 min, and filtered through 0.45-μm PVDF filters. The eVLPs were then concentrated by ultracentrifugation at 50,000 × g for 2 hours at 4°C and resuspended in 100 μl cold PBS.

For eVLP transduction, organoids were isolated from BME, enzymatically digested using TrypLE Express Enzyme for 3 min at 37°C to make single cells, and resuspended in expansion medium supplemented with 8 μg/ml Polybrene (Sigma, TR-1003-G) and 10 μM Y-27632. The cells were then transferred to 48 well suspension plates in a volume of 100 μl per well and mixed with 25 μl eVLPs. The plate was centrifuged at 500 × g for 1 hour at 32°C and then incubated for 3 hours at 37°C. Afterwards, the cells were isolated, plated on 24 well plates in BME droplets, and grown in expansion medium supplemented with 10 μM Y-27632 until organoid structure was formed.

### Genotyping

Genomic DNA was isolated using Quick DNA microprep kit (Zymo Research, D3021) according to the manufacturer’s instructions. The region of interest in the *HBEGF* locus was amplified using Q5 Hot Start High-Fidelity 2x Master Mix (NEB, M0494) with 32 cycles of 98°C for 10 sec, 59°C for 30 sec, and 72°C for 30 sec. PCR products were purified using NucleoSpin Gel and PCR Clean-Up Kit (Macherey-Nagel, 740609.50). Purified PCR products sent for Sanger sequencing using the forward primer at Macrogen Europe. PCR primers can be found in Supplementary Table 2. Gene editing efficiency was assessed using EditR web application (Kluesner, 2018).

### Synthetic hydrogel fabrication and encapsulation

Organoids were mixed with appropriate volumes of 25 wt% polyethylene glycol with vinyl sulfone functional group (PEG-VS; 4-arm 20 kDa; JenKem) precursor solutions and stoichiometrically balanced ratios of reactive groups in di-thiol MMP-degradable peptide (KCGPQGIWGQCK; Genscript) and RGD peptides (CRGDS; Genscript) to generate hydrogel networks of a desired final PEG content (3-4 wt%).

For 3 wt% final concentration PEG hydrogels the organoids were mixed with sterile solution consisting of a PEG-VS (3 wt%; VS = 6 mM), MMP-degradable peptide (SH = 5 mM) and RGD peptides (SH = 1 mM). The pH for all solutions was adjusted to 7.4. Gels were allowed to crosslink by incubating them at 37 °C for a minimum period of 30 min or until the hydrogels solidified. After which medium was added to hydrogel-encapsulated organoids.

### Statistical analysis

Statistical mean comparison tests were performed in GraphPad Prism 9. Comparisons between 2 conditions were performed using unpaired t-test or ratio paired t-test if multiple matched donors were compared. Comparisons between 3 or more conditions were performed using one-way ANOVA followed by Tukey’s multiple comparison test. A p-value less than 0.05 was considered to be statistically significant. Sample size varied for each experiment and indicated in the figure legends. Biological replicates were defined as independent organoid/cell cultures (e.g., different wells/plates). Unless stated otherwise, bar graphs with linear scale are shown as mean ± standard deviation, bar graphs with logarithmic scale are shown as mean ± standard error of the mean, box-and-whisker plots are shown as median (line), interquartile range (box), and data range or 1.5x interquartile range (whisker).

## Acknowledgments

The study was supported by ZonMW TAS (‘Regenerating Intestinal Tissue with Stem cells’ project), Metakids (‘Minilevertjes’ project) and the ERC starting grant funding (all to S.A.F.).

