## Supplementary Figures for "Novel hepatocyte-like liver organoids recapitulate crucial mature hepatic functions"

**Supplemental Figures:**

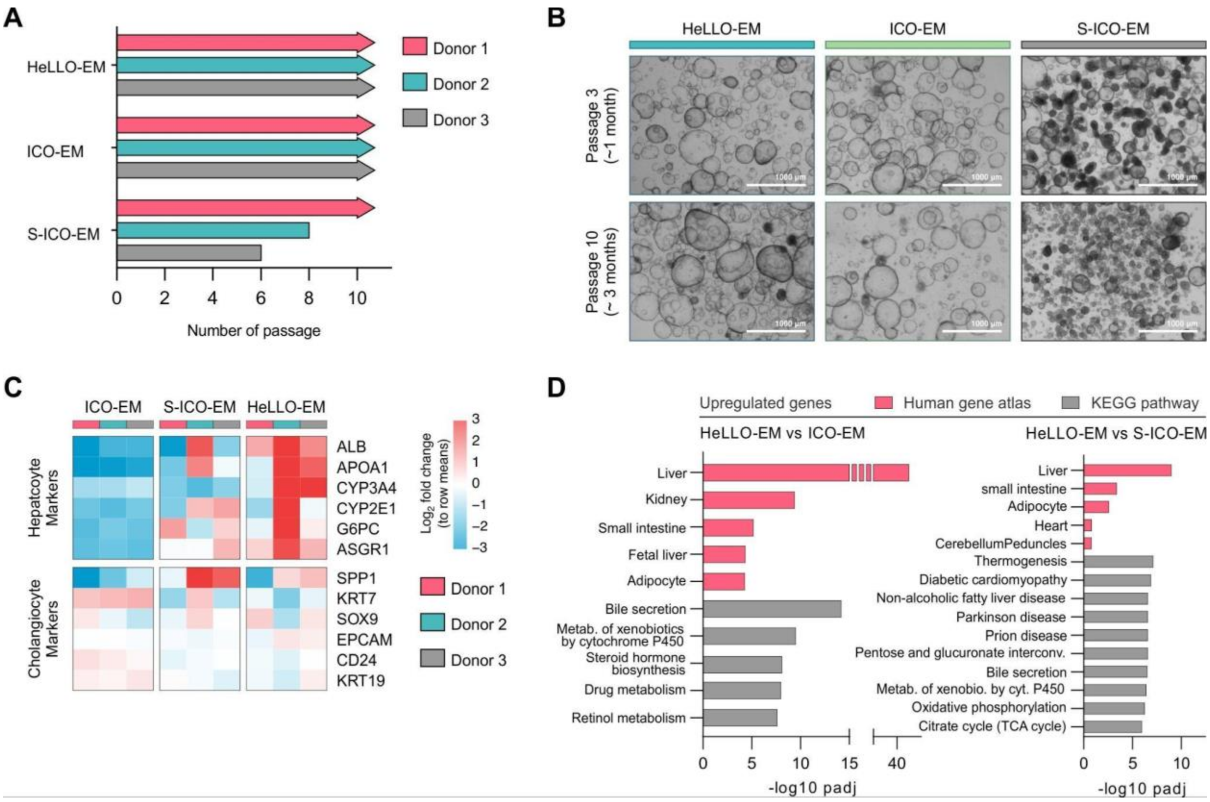


**Figure S1: HeLLO-EM is more hepatic than ICO-EM.**

**(A)** Passaging of organoids from three donors cultured with HeLLO expansion medium (HeLLO-EM), Huch ICO expansion medium (ICO-EM) and Sampaziotis ICO expansion medium (S-ICO-EM) showing similar expansion capacity between HeLLOs and ICOs. **(B)** Brightfield images of donor-matched HeLLOs, ICOs and S-ICOs in expansion medium at passage 3 and passage 10. **(C)** Transcriptomic analysis of donor-matched HeLLOs, ICOs and S-ICOs in expansion medium in passage 3 shows much higher hepatic expression and lower cholangiocyte expression in HeLLOs. **(D)** Enrichment analysis using differentially expressed genes shows a major increase in liver gene expression in HeLLO-EM based on human gene atlas and KEGG pathway databases.

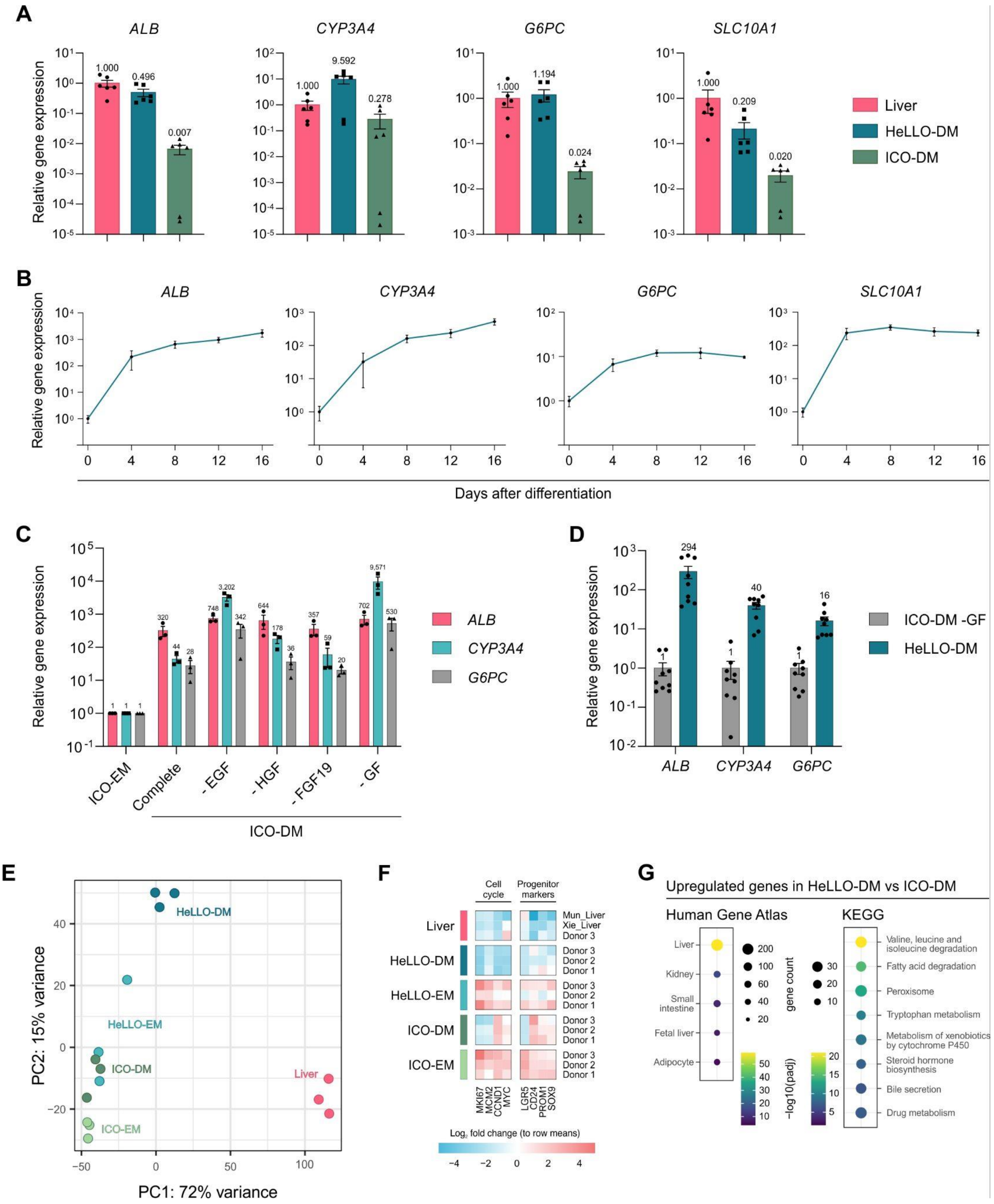

**Figure S2: Investigating the importance of the medium factors for the HeLLO hepatocyte phenotype. HeLLO-DM displays stronger hepatic resemblance due to the combination of optimized expansion medium and removal of growth factors in differentiation medium.**

**(A)** Quantitative PCR analysis of four hepatic marker genes for liver tissue, HeLLO-DM and ICO-DM (data from three donors and two biological replicates). **(B)** Quantitative PCR analysis of four hepatic marker genes for HeLLO-DM over time from the moment of switching from EM to DM. **(C)** Quantitative PCR analysis showed that removal of several growth factors (GFs) in ICO-DM increased the expression of hepatocyte marker genes (data from three biological replicates from two donors). **(D)** Quantitative PCR analysis of three hepatic marker genes for ICOs differentiated in ICO-DM without growth factors

(ICO-DM -GF) and HeLLOs differentiated in HeLLO-DM showing that expansion medium composition is also important in generating a strong hepatic phenotype in differentiation medium (data from 9 biological replicates from 2 donors). **(E)** PCA analysis of donor matched ICOs and HeLLOs in expansion and differentiation media (day 8) and liver (two extra liver samples were integrated using HLCompR). Transcriptomic data showed a shift towards a more liver phenotype of HeLLO-DM. **(F)** Heatmap analysis showing a strong decrease of cell cycle and progenitor marker expression levels in HeLLO-DM. **(G)** Enrichment analysis using differentially expressed genes showed a major increase in liver gene expression in HeLLO-DM compared to ICO-DM based on human gene atlas and KEGG pathway databases (data from three donors).

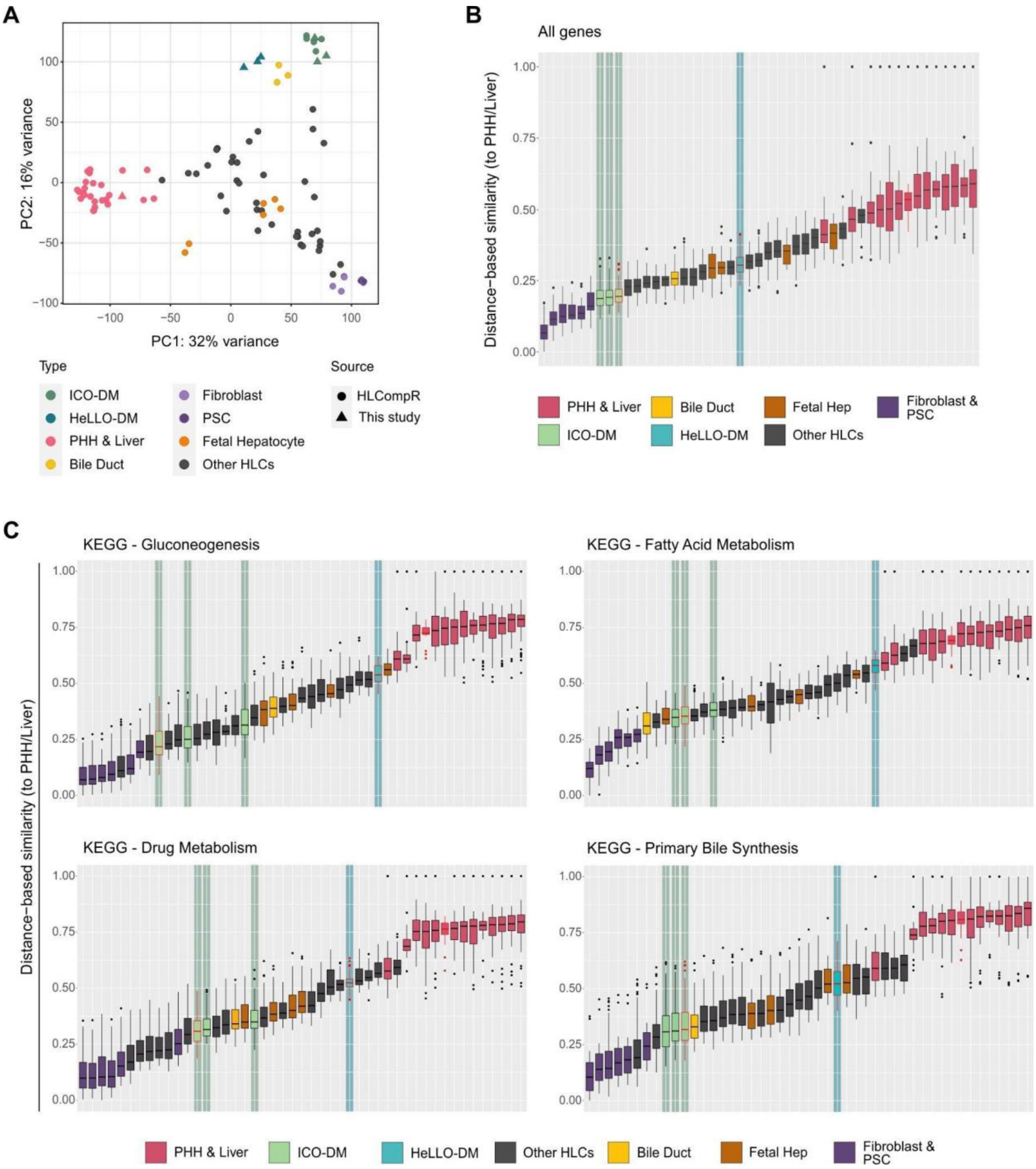


**Figure S3: HeLLOs strongly express important hepatic pathways compared to other hepatocyte models.**

(A) PCA analysis generated by HLCompR showing an increase of hepatic phenotype in HeLLO-DM when compared to ICO-DM and other hepatocyte models. (B) Euclidean distance based similarity generated by HLCompR showing that HeLLO-DM (blue highlight) is in overall more similar to PHH/liver than ICO-DM (green highlight). (C) Euclidean distance based similarity generated by HLCompR showing that HeLLO-DM excels at specific hepatic pathways compared to liver and other hepatocyte models.

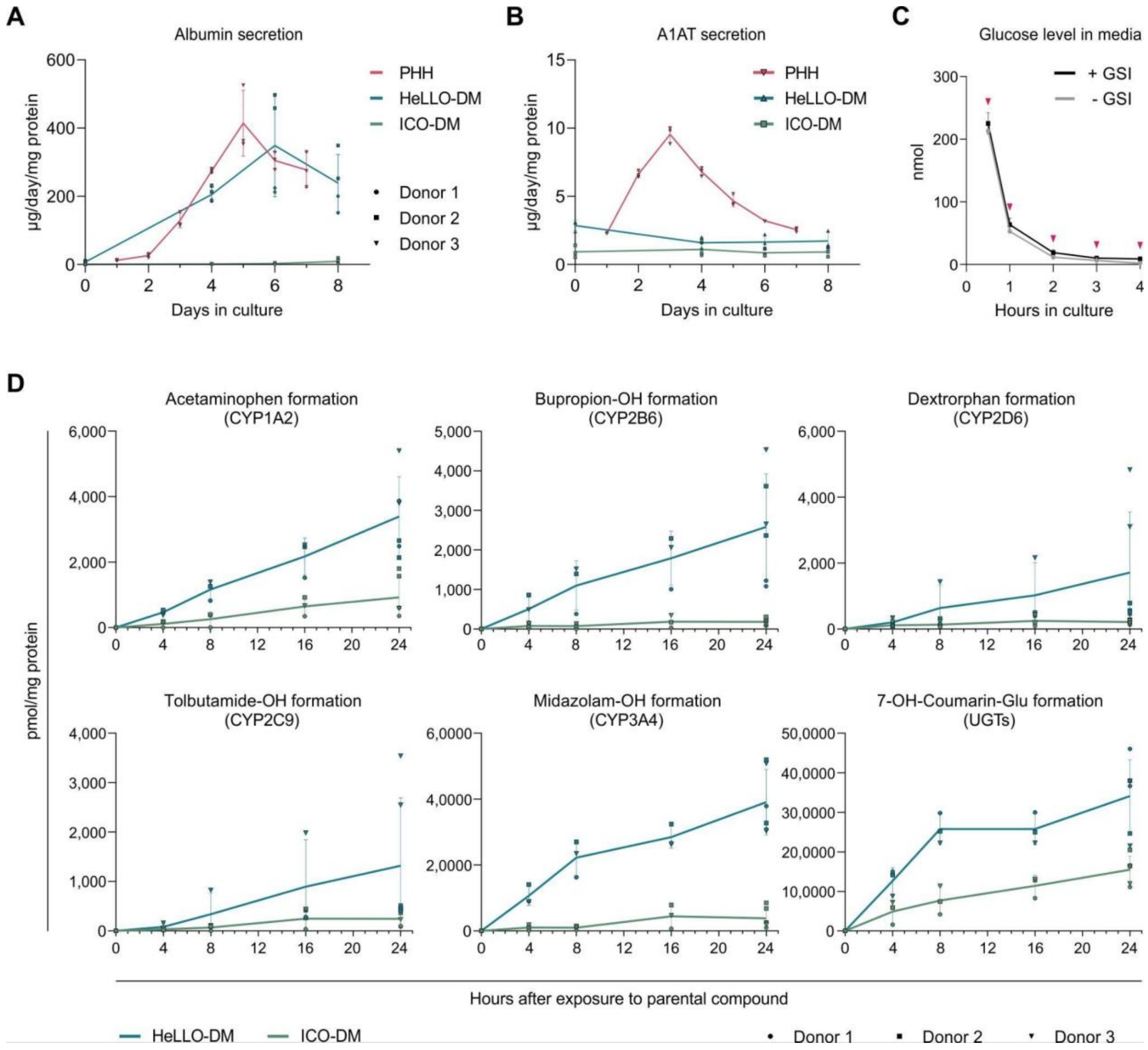


**Figure S4: Time course analysis of hepatic functions in HeLLO-DM and ICO-DM.**
**(A)** Albumin secretion as measured by ELISA for sandwich-cultured PHHs, HeLLO-DM and ICO-DM showing similar albumin production of HeLLO-DM compared to PHHs (data from two matched donors for HeLLO-DM and ICO-DM, three biological replicates from one donor for PHHs). **(B)** A1AT secretion as measured by ELISA for sandwich-cultured PHHs, HeLLO-DM and ICO-DM showing a general increase of A1AT production for HeLLO-DM compared to ICO-DM (data from two matched donors for HeLLO-DM and ICO-DM, three biological replicates from one donor for PHHs). **(C)** Glucose measurement of glucose-free differentiation medium after serial refreshment (arrowheads) on HeLLO-DM (data from three biological replicates from one donor). **(D)** Drug metabolism activity for HeLLO-DM and ICO-DM for several substrates over time showing much higher activity in HeLLO-DM (data from three matched donors).

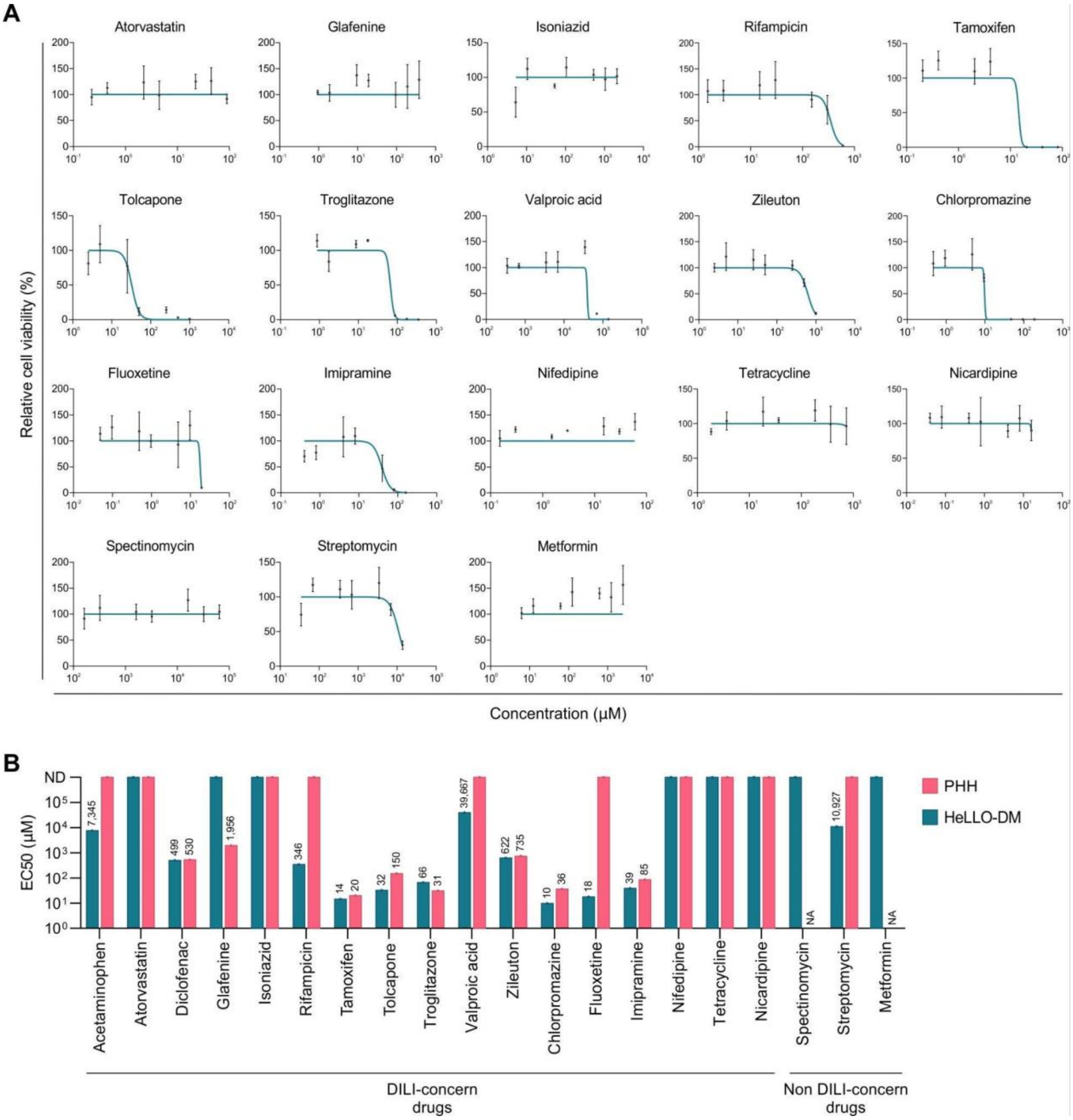


**Figure S5: HeLLO-based toxicity screen effectively predicts toxicity of various known liver-toxic drugs.**

**(A)** Toxicity response of HeLLOs to 15 DILI-concern drugs as represented in Fig. 4C. Spectinomycin, streptomycin, and metformin (non DILI-concern drugs) were included as negative controls (data from one donor, three biological replicates). **(B)** EC50 values of all drugs tested on HeLLOs (data from one donor, three biological replicates) and 2D cultured PHHs (data from Li et al. 2020).

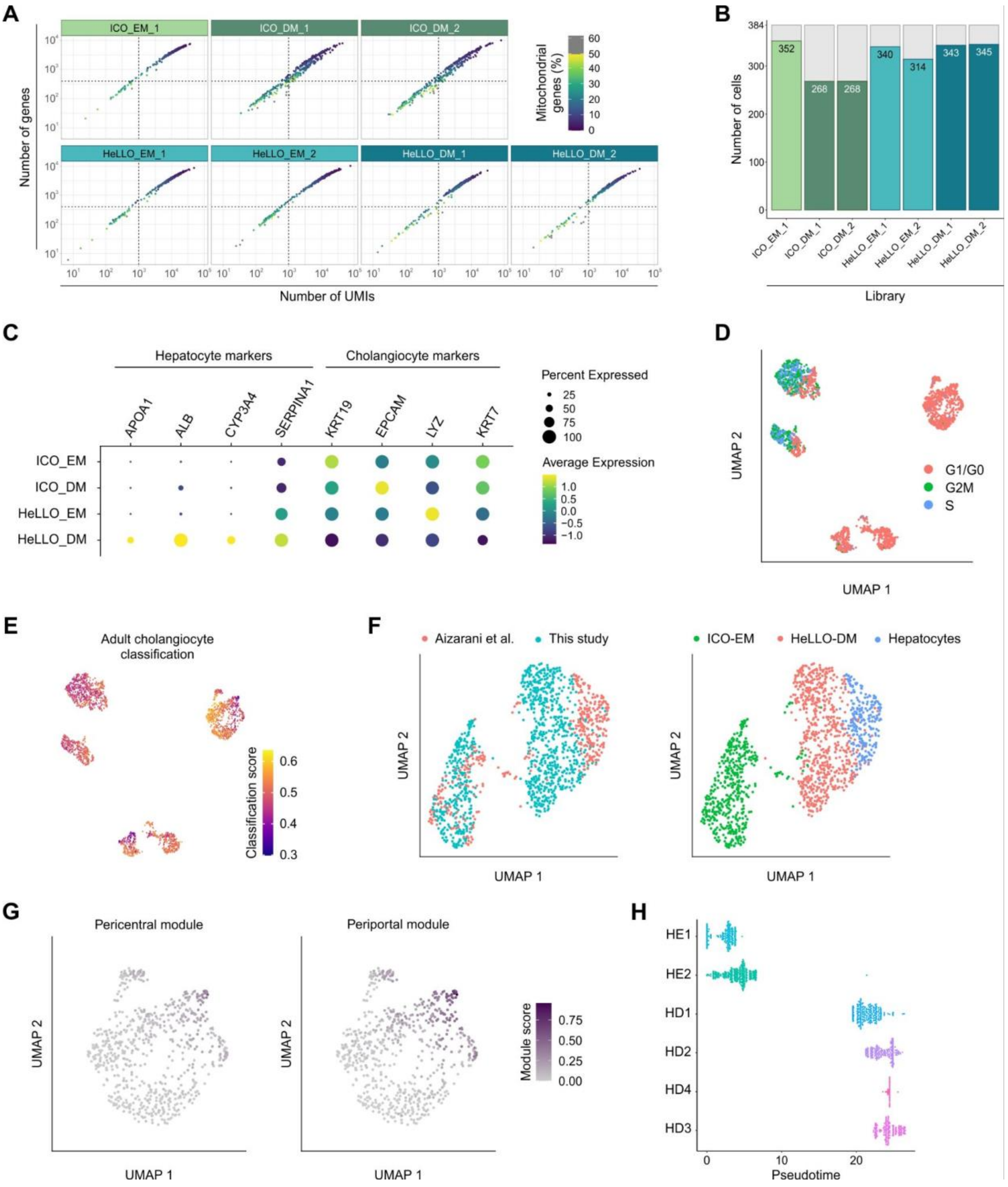


**Figure S6: Single-cell transcriptomic analysis of HeLLO-EM and HeLLO-DM uncovers heterogeneity within the HeLLO culture.**

**(A)** Scatter plot showing the number of genes and unique molecular identifiers (UMIs) in each sample library. Color indicates the percentage of mitochondrial genes and dashed line represents the threshold used to filter out low quality cells (number of genes = 400; number of UMIs = 1000). **(B)** Bar graph showing the number of cells retained after filtering out low quality cells per sample library. Gray color indicates the number of cells before filtering (384 cells). **(C)** Dot plots of important hepatic and cholangiocyte marker expression level (color) and percentage of cells expressing the marker gene (dot size) in each organoid type. **(D)** UMAP visualization of the cell cycle classification based on cell cycle gene scoring. **(E)** UMAP showing SingleR classification scores for adult cholangiocytes from the human liver development atlas (Wesley, 2022). **(F)** UMAP analysis of integrated single-cell sequencing data of this study and Aizarani et al. (2019). Annotations of the study and sample type are shown on the left and right graphs respectively. ICO-EM from both studies clustering together indicates a proper integration. **(G)** UMAP showing the module score for pericentral and periportal gene sets. **(H)** The distribution of cells on the pseudotime from each HeLLO cluster showing that HD1 arises earliest in pseudotime while HD3 arises latest among the other HeLLO DM clusters.

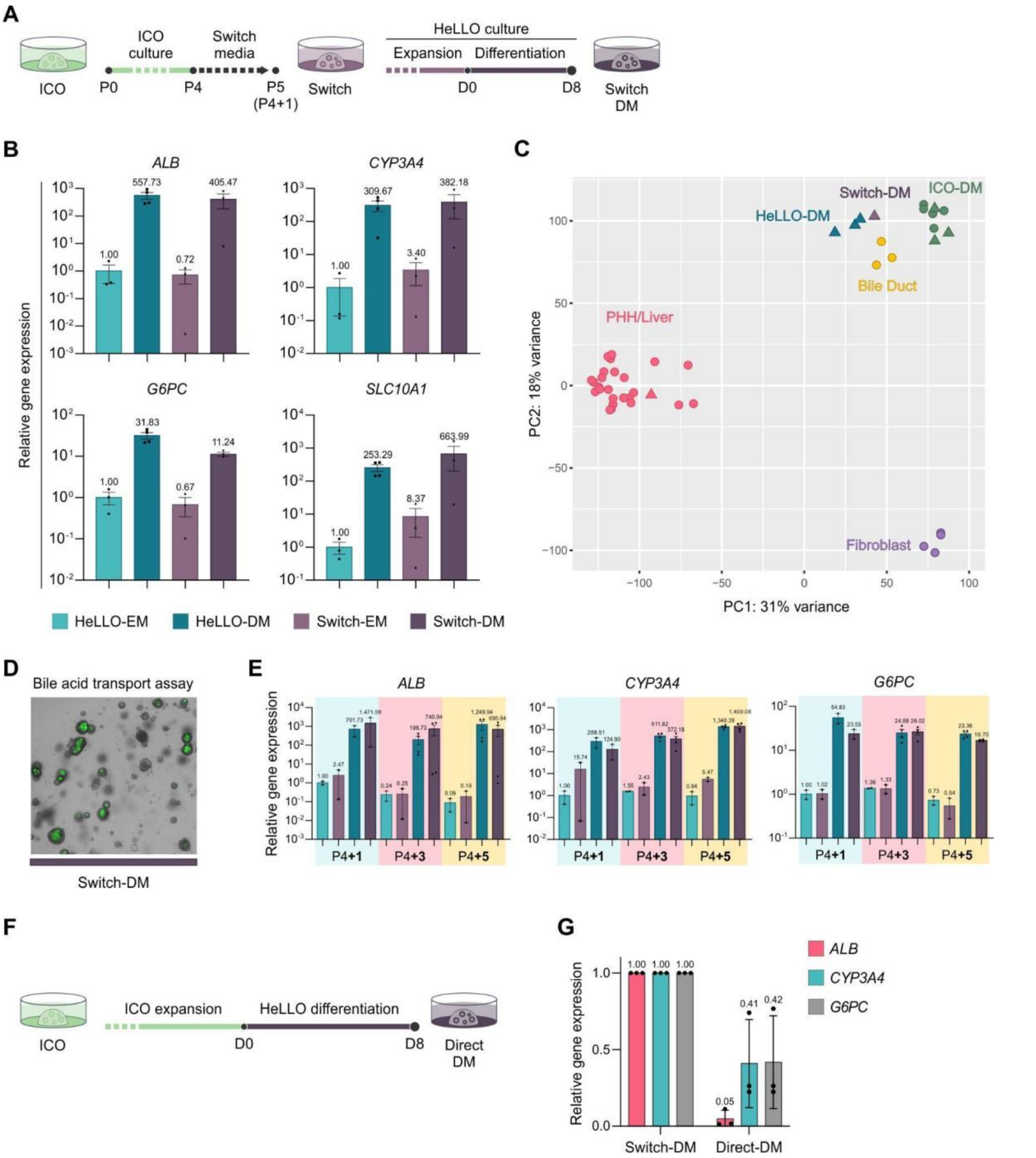


**Figure S7: ICOs can be differentiated into HeLLO-resembling cultures to improve hepatic differentiation.**

**(A)** Illustration of the switch condition, which starts from ICO-EM culture and then switches to HeLLO-EM and HeLLO-DM culture. **(B)** qPCR analysis indicates a strong induction of hepatic differentiation in the switch condition similar to organoids established and grown as HeLLOs (data from three biological replicates). **(C)** PCA analysis of transcriptome data shows an improvement in hepatic differentiation of the switch condition compared to ICOs, while HeLLOs remain the most hepatocyte-like (data from one donor). **(D)** The switch condition enables bile acid transport assay which the ICOs are incapable of facilitating. **(E)** qPCR analysis of different differentiation timings in the switch condition indicating stable improvement of hepatocyte differentiation (data from three biological replicates). After switching at passage 5, the organoids were differentiated at passage 5 (P4+1), passage 7 (P4+3), and passage 9 (P4+5). **(F)** Illustration of using HeLLO-DM on the ICOs directly after expanding. **(G)** qPCR analysis of directly adding HeLLO-DM to ICOs showing a decrease in hepatocyte differentiation compared to the switch condition (data from three biological replicates).

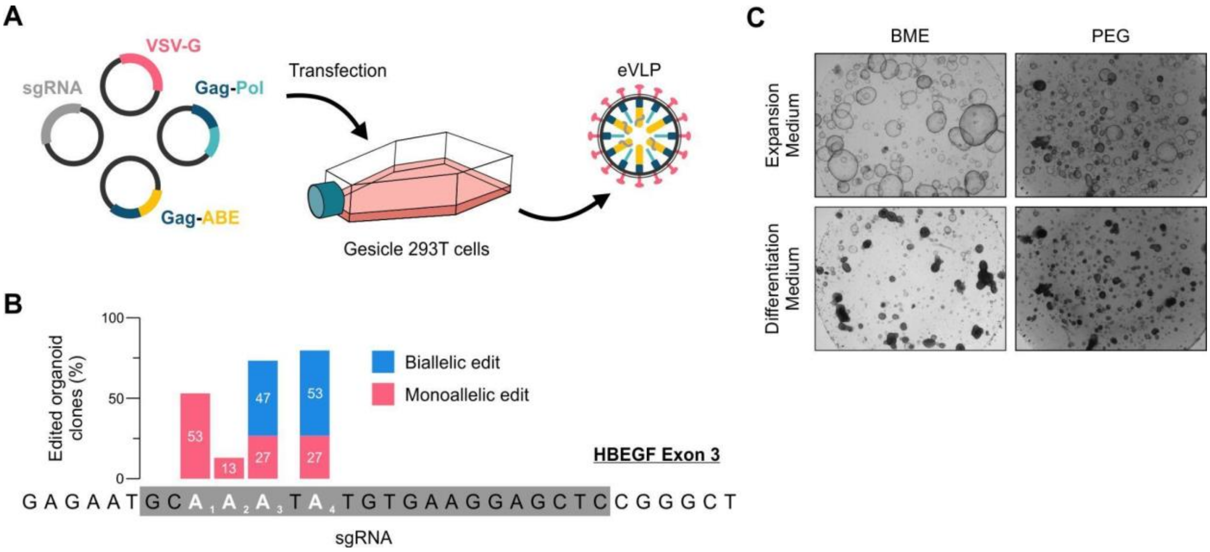


**Figure S8: Applying novel labtools on the HeLLOs.**
**(A)** Schematic overview of the production method of base editing eVLPs (Banskota, 2022). **(B)** HeLLOs treated with eVLPs targeting HBEGF exon 3 shows high editing signal based on sanger sequencing. **(C)** HeLLOs cultured in a 4% polyethylene glycol (PEG) synthetic hydrogel supplemented with arginylglycylaspartic acid (RGD) peptides were viable and similar to matrigel/BME.
